# A near-ubiquitous plasmid characterises Second Pandemic *Yersinia pestis*

**DOI:** 10.64898/2026.10.03.756212

**Authors:** Frederik V. Seersholm, Gabriele Scorrano, Jon Bang Ploug, Ruairidh McLeod, Sam E. Williams, Marie Louise S. Jørkov, Lutz Klassen, Stig Bergmann Møller, Djillali Hadjouis, Esben Klinker Hansen, Mikkel B. Siebken, Jane Jark Clausen, Stine Damsbo Winther, Mads Ravn, Caroline Ahlström Arcini, Astrid K.N. Iversen, Eske Willerslev, Martin Sikora

**Affiliations:** Centre for Ancient Environmental Genomics, Globe Institute, University of Copenhagen, Copenhagen, Denmark; Center for Molecular Anthropology for the study of ancient DNA, Department of Biology, University of Rome Tor Vergata, Rome, Italy; PandemiX Center, Department of Science and Environment, Roskilde University, Denmark; Department of Communication and Arts, Roskilde University, Denmark; All Souls College, University of Oxford, Oxford, UK; ArchaeoScience, Section for Geogenetics, Globe Institute, University of Copenhagen, Copenhagen, Denmark; Museum Østjylland, Stemannsgade 2, Randers, Denmark; Arkæologi Nordjyske Museer, Aalborg, Denmark; Laboratoire départemental d’archéologie du Val-de-Marne, 3, rue Guy-Môquet, 94800 Villejuif, France; VejleMuseerne, Vejle, Denmark; Museum of Copenhagen, Copenhagen, Denmark; Department of Local Heritage, Moesgaard Museum, Aarhus, Denmark; Department of Clinical Sciences, Lund University, Lund, Sweden; Nuffield Department of Clinical Neurosciences, John Radcliffe Hospital, Weatherall Institute of Molecular Medicine, University of Oxford, Oxford, UK; Department of Genetics, University of Cambridge, Cambridge, UK

## Abstract

*Yersinia pestis*, the causative agent of plague, crippled population across Eurasia in three well-described historical pandemics: The Justinianic plague (541-780), the Second Pandemic (1346-1840) and the Third Pandemic (1894-). The Second Pandemic was the deadliest, with the Black Death (1346-1351) killing up to 60% of Europe’s population, far exceeding the population mortality of the Third pandemic, despite few genomic differences between their plague strains. To identify unique genetic variation that could help explain the high death toll of the Second Plague pandemic, we used variation-graph based pan-genomics on a large dataset of 110 new and 12 published shotgun sequenced ancient plague genomes to comprehensively characterise the pangenome diversity of the Second Pandemic plague. Our pangenomic approach revealed the presence of a cryptic 34kb plasmid, pTP33, in almost all strains from the Second Pandemic, but undetectable in all other ancient strains. We find that pTP33 is a phage-plasmid-like derived element with homologues in Enterobacterales bacteria with established insect- or invertebrate-associated lifestyle phases. Our results suggest that pTP33 widened the range and infectivity of vector species available for plague transmission, consistent with reported effects on biofilm formation and bacterial retention. We hypothesise that pTP33 broadened the vector range of *Y. pestis* beyond the classical rat/flea cycle, facilitating the establishment of new rodent-ectoparasite plague reservoirs, which, critically, increased the risk of flea- or lice-mediated plague transmission between humans. Taken together, our findings offer a biological explanation for differences in transmission dynamics and population mortality between the Second and Third plague pandemics.

## Introduction

The Second Plague Pandemic is one of the deadliest disease outbreaks known to humanity, and its dramatic socio-economic consequences shaped the course of history. The first wave of the pandemic, known as the Black Death, originated in central Asia around 1339 AD^1^ from where it swept across Europe (1346-1353 AD) killing an estimated 60% of the European population^2^. Over the course of the coming 500 years, the disease would propagate through Eurasia in epidemic waves, costing the lives of hundreds of million people globally^2^.

Because of its devastating consequences, the Black Death, and the subsequent Second Pandemic outbreaks are among the most extensively studied historical pandemics, with insights obtained from a wide array of different sources, such as historical records^3^, pollen^4^, bioarchaeology^5^, human selection patterns^6^ and ancient pathogen DNA^7^. For example, palynological studies have demonstrated that the impact of the Black Death was so severe that the widespread abandonment of farmland caused forest regrowth of such magnitude that it is visible in the pollen record^4,8^.

The causative agent of plague, the bacterium *Y. pestis*, was one of the first pathogens to be studied when the field of ancient DNA was in its infancy^9^, and the full genome of the bacterium was already published in 2011^10^. The ensuing scientific publications studying the diversity within the Second Pandemic have demonstrated diversification within the bacterium, with some geographical and temporal structure, in which specific strains can be linked to well described outbreaks of the disease^11^. Surprisingly, there is currently very little evidence for large genomic differences between the strains of the Second Pandemic and the later Third Pandemic. In fact, phylogenetic reconstructions of the chromosome of ancient and modern plague have found that the Second Pandemic plague strains fall within the modern diversity of plague on branch 1 of the plague phylogeny^11^.

Given the low chromosomal diversity between Third and Second Pandemic plague strains, it has previously been suggested that the difference in disease manifestation and virulence might be explained by variations in plasmid content of the bacterium^12^. *Y. pestis* has three almost ubiquitous and well described plasmids: pCD1 (70kb), pPCP1 (9.6kb), and pMT1 (96kb), of which the latter two are found exclusively in *Y.pestis*, whereas pCD1 are present in *Y.pseudotuberculosis* and *Y. enterolitica* alike. The role of these three plasmids in plague virulence is well established. The pPCP1 plasmid, for example, carries the *pla* gene that is associated with high disease virulence in humans and rats. It was recently hypothesised that *pla* gene depletion in late Second Pandemic strains resulted from selection ffor attenuatedvirulence following the massive plague-induced reduction and fragmentation of rat populations to allow for persistence and continued transmission of plague^13^. Similarly, the *ymt* gene on the pMT1 plasmid is associated with survival of the bacterium in the midgut of fleas, and accordingly the pathogens ability to cause bubonic plague^14^. Lastly, the type III secretion system (T3SS) encoded on pCD1 produces *Yersinia* outer proteins (Yops) that inhibits host immune function in *Y.pestis*, *Y.pseudotuberculosis* and *Y. enterolitica*^15,16^. The progressive loss and gain of some genes within these plasmids have been proposed, for example *ymt* has been shown to have been obtained immediately after the LNBA- and LNBA+ clade divergence approximately 3800 years BP^17^. Although it is well-established that the acquisition of plasmids can provide bacteria with beneficial traits that facilitate adaptation to changing environmental conditions and/or transmission routes^18^, the role of cryptic or rare accessory plasmids in ancient plague samples has not been studied, and the precise genetic composition of ancient plague plasmids remains to be described in a pangenomic framework.

In order to identify genomic elements associated with the devastating effects of the Second Pandemic, we generated shotgun sequencing data of 110 new plague positive shotgun sequenced samples from the Second Pandemic, and characterised their pangenomic diversity together with already published plague genomes using a variant-graph based approach.

## Results

### 110 new shotgun sequenced plague samples

We screened 1,071 recently sequenced human samples from the Middle Ages for presence of *Y. pestis* DNA and identified a total of 110 plague positive samples (Supplementary Table 1). The samples range in plague coverage from 0.0013X to 23.4X (median: 0.028), with a total of six samples over 4X which we use as the threshold for traditional SNP-calling and phylogenetic analysis. By merging these samples with a panel of previously published plague genomes, we generate a final data set of 228 plague cases from the Second Plague Pandemic, spread across most of Europe (see Fig. 1a) and dated to between 1346 and the beginning of the 19th century. The vast majority (106/118; 90%) of previously published plague genomes were enriched for plague DNA using targeted-capture approaches. All the samples presented in this study, on the other hand, were generated using either shotgun sequencing exclusively, or a combination of shotgun sequencing and capture. As such, we increase the volume of available shotgun plague data from the Second Pandemic tenfold, from 12 previously published genomes to 122 (see Fig. 1b).

**Figure 1.**
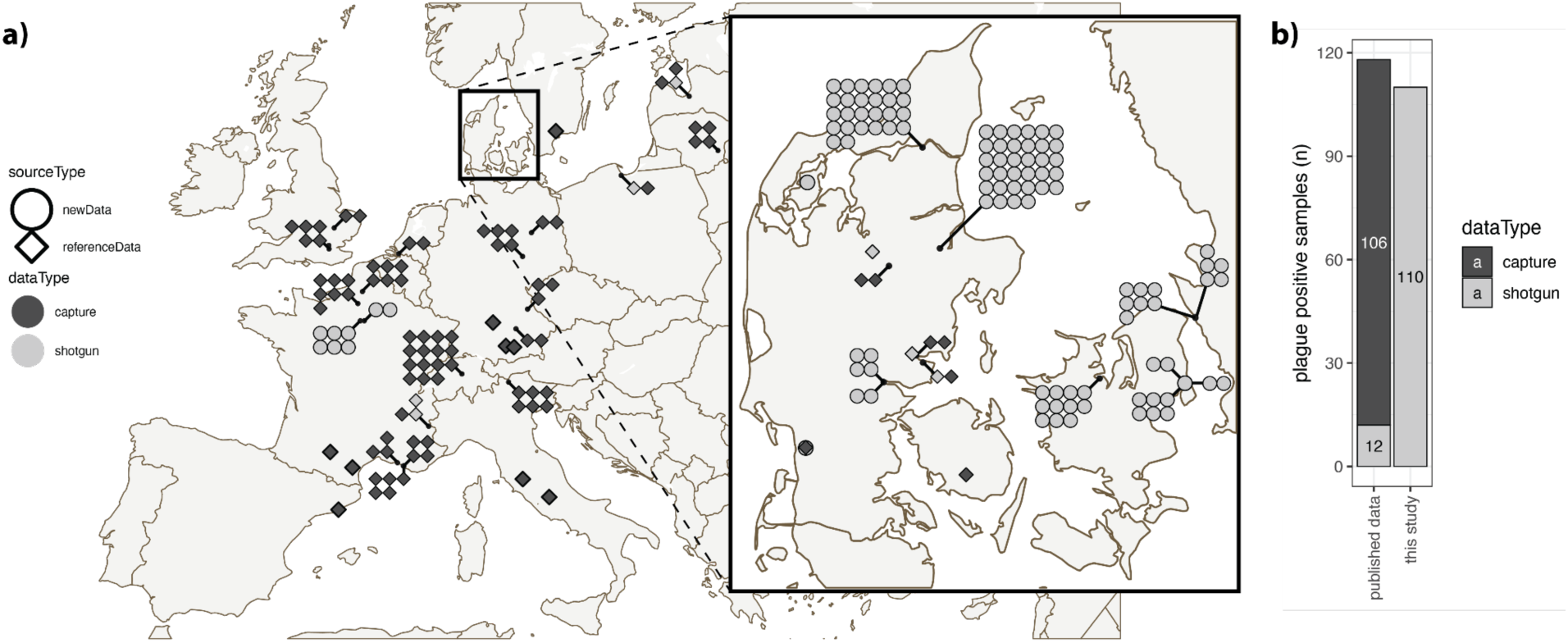
Overview of Second Pandemic plague samples. **a)** Map of plague positive samples dated between 1346 and 1800, with sequencing strategy indicated by color. New samples published in this study are round shapes while diamond shapes represent previously published samples. **b)** Barplot comparing the number of plague samples already published with the number of samples presented in this study.

Reflecting the geographical distribution of the human dataset screened, the vast majority (n=90) of our samples are from Denmark, with an additional 12 and 8 new samples from Sweden and France, respectively. Importantly, we present the two single sites with the highest number of plague infected individuals published so far: Tjærby, Denmark with 34 infected individuals and Vor Frue Kirkegård, Denmark with 26 infected individuals.

### Phylogenetic placements

To investigate the phylogenetic structure within our dataset, we reconstructed a phylogenetic tree using all plague genomes with a coverage over 4X (Fig. 2a, Supplementary Fig. 1, Supplementary Fig. 2). We classified the phylogeny into five manually assigned clades from the most basal to the most derived: basal1, basal2, middle, preLate and late. We added a total of six new samples with a plague coverage over 4X to this phylogeny: One sample from Sweden (CGG105888; Saint Cllemens) belongs to the clade of the Black Death genomes in ‘basal1’, while two genomes from Tjærby, Denmark (CGG100891 and CGG100920), together with CGG100425 from Vor Frue Kirkegård falls within the “basal2” clade. Lastly, we find that samples CGG100462 and CGG100422 from Vor Frue Kirkegård cluster within clade “middle” (Fig. 2a, Supplementary Table 1, Supplementary Table 2).

**Figure 2.**
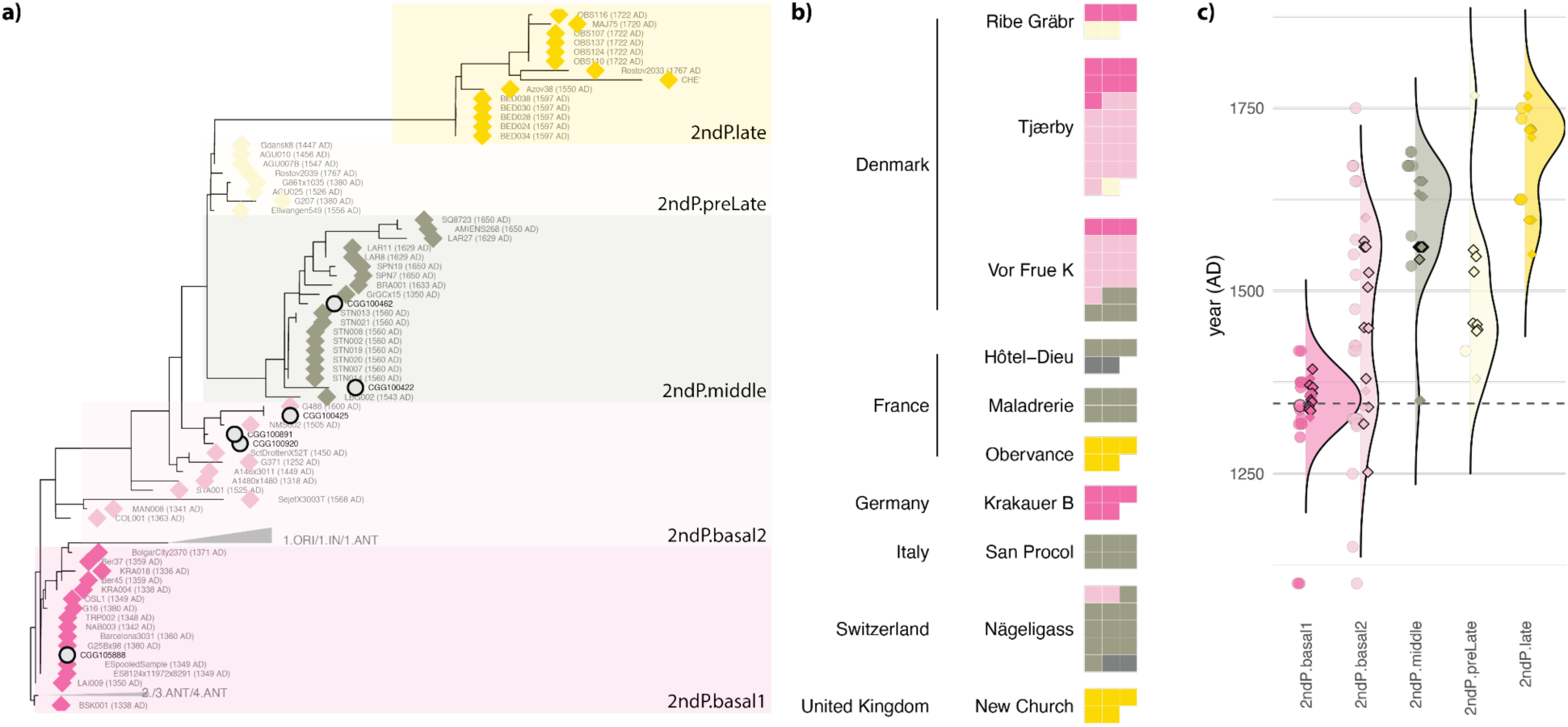
Phylogenetic context of all samples analysed. **a)** Phylogenetic tree of all high coverage samples (>4x). **b)** Overview of phylogenetic placements of both high and low coverage (>0.01X) samples from sites with more than five plague cases. **c)** Age distribution of strains within each plague clade with density distributions calculated using dates from all reference genomes and ages from all new strains with a radiocarbon date. Diamonds and circles represent reference data and data from this study respectively, while a black outline highlights radiocarbon dated strains.

To also characterise lower coverage genomes phylogenetically, we assigned all genomes with coverage between 0.01X and 4X (n=116) to one of the five subclades described above based on phylogenetic placement analysis (see Fig. 2b, Supplementary Fig. 3, Supplementary Fig. 4, Supplementary Fig. 5, Supplementary Fig. 6, Supplementary Fig. 7, Supplementary Fig. 8). Using these approaches, we were able to classify 74% (169/228) of all samples in the database (reference genomes and new samples), and assign these to a subclade. We found that most of the largest sites studied had evidence of multiple plague outbreaks during the pandemic as distinct lineages from different subclades were often identified within single sites. For example, we identified plague strains from lineages ‘basal1’, ‘basal2’ and ‘middle’ within Vor Frue Kirkegård and strains from ‘basal1’, ‘basal2’ and ‘preLate’ in Tjærby Fig. 2. Generally, we find that the ages of samples from the more basal clades are younger than that of the derived ones (Fig. 2c), however, there is a relatively large spread within each clade. This observation is most likely attributable to the uncertainties associated with radiocarbon dating of samples from the Middle Ages^19^. However, it is also expected to find some variation in age within these clades. For example, even though the genomes G488 and CGG100425 belong to the clade ‘basal2’ they fall on a long-derived branch within this clade, and accordingly their ages are markedly younger than the rest of the clade.

### Plasmid diversity during the Second Plague Pandemic

To characterise Second Pandemic genetic diversity beyond the canonical plasmids and the chromosome sequence of the standard *Y. pestis* reference genome (see Methods), we downloaded all published complete assemblies within the *Y. pseudotuberculosis* species complex to create a pangenome variation graph. In total, the input sequences for the graph building consists of 698 plasmid sequences and 275 chromosomal sequences from 246 assemblies, with 223 assemblies from *Y. pestis*, 21 from *Y. pseudotuberculosis* and two from *Y. similis*. Unsurprisingly, we find plasmids pCD1, pMT1 and pPCP1 to be almost ubiquitous in modern *Y.pestis* assemblies, while only pCD1 is consistently found in *Y. pseudotuberculosis* assemblies (Fig. 3a, Supplementary Table 3). For *Y. similis*, we found no consistent pattern with one distinct plasmid found in each assembly (Fig. 3a).

**Figure 3.**
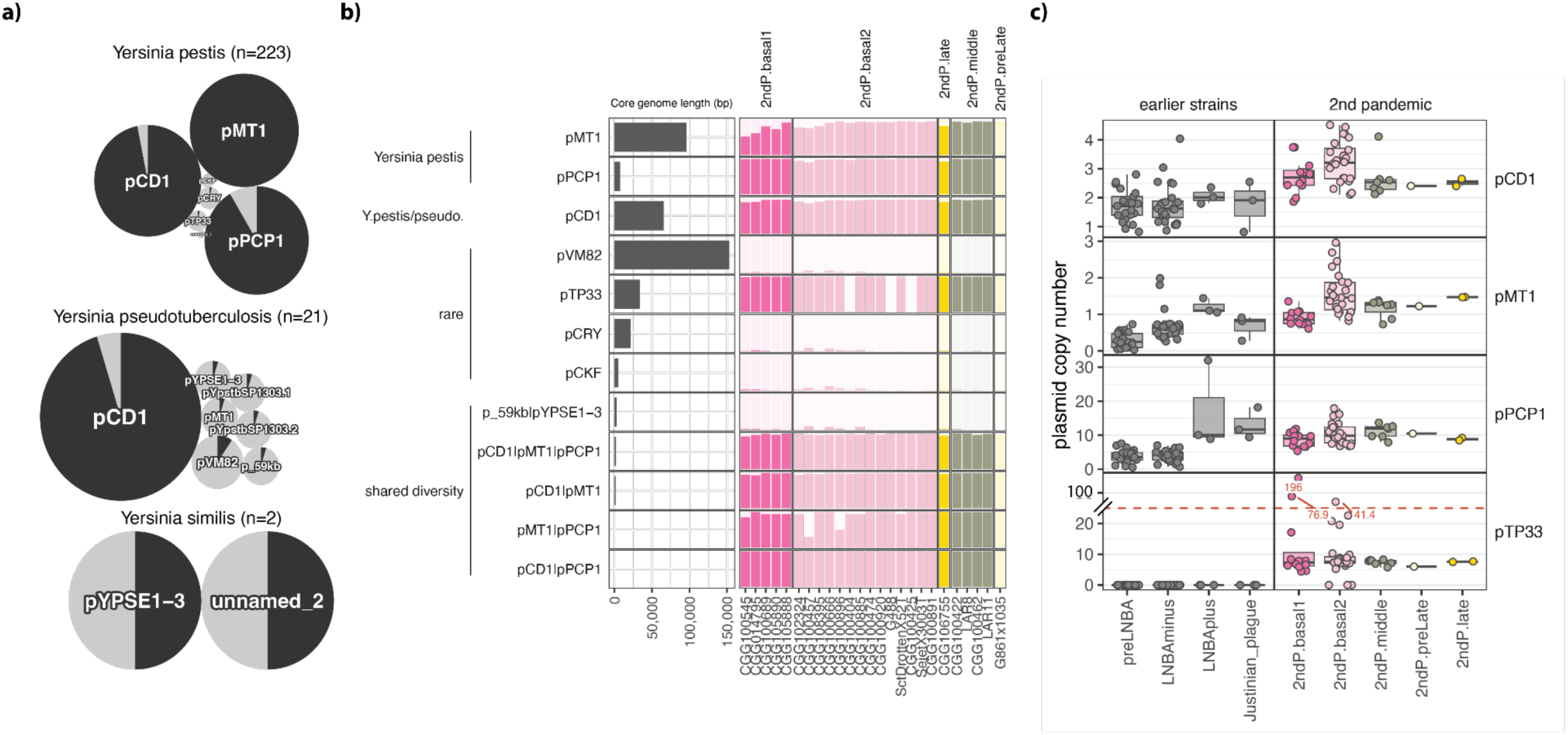
Overview of pangenome analysis. **a)** Plasmid presence/absence for all assemblies used to build the pangenome variation graph. Pies are scaled by plasmid frequency in each species (Supplementary Table 3). **b)** Ancient DNA mapping results to the pangenome variation graph. Left side panel: total length of the core genome for each category. Categories represent either single plasmids or multiple plasmids in cases where genetic nodes are present in multiple plasmids (shared diversity). Right side panel: Fraction of core genome with coverage for each shotgun sample (>1X) stratified by plasmid category. **c)** Plasmid copy number estimates calculated as the average coverage of core genome nodes from the pangenome for each plasmid normalised to the coverage of the chromosome. Because of three outliers with very high copy number from the pTP33 plasmid, the y-axis has been truncated for this part of the plot (red stippled line).

Using these assemblies, we built pangenome variation graphs for the plasmids and the chromosomes. The final plasmid graph consists of 13,399 nodes spanning 1,366,414 bp, of which 483,053 bp (35%) are observed in more than one assembly. Focusing on the core genome of each plasmid community, we find that the total graph sizes for each plasmid are similar but slightly shorter than the canonical plasmid lengths from the literature: pMT1=90.9 kb (refseq: 96.2 kb), pPCP1=7.6 kb (refseq: 9.6 kb) and pCD1=67.9 kb (refseq: 70.2 kb)^20^ (Fig. 3b, left side panel). This length discrepancy is explained in large part by a variable number of repeated insertion sequences (IS elements, IS100, IS285-derived elements (pMT1 and pCD1), and IS1541 (pMT1)); mobile genetic elements that can promote gene rearrangements and are grouped as ‘shared diversity’ (Fig. 3b, Supplementary Fig. 9).

As expected, the mapping of ancient shotgun data to the graph, revealed that the three canonical *Y.pestis* plasmids were present in all ancient samples (Fig. 3b, right side panel) and that the majority of genetic diversity in the chromosome is shared among all three species of the *Y. pseudotuberculosis* species complex (Supplementary Fig. 10). For the three canonical plasmids, we generally find that very close to 100% of the core genome variation from the graph is present in all samples, with a slight drop-off noted for the low copy number plasmid pMT1 in samples with lower coverage (left most samples within each subclade). Similarly, we also find that the diversity shared between these canonical plasmids, such as insertion sequences (e.g., IS100) is detectable in all the ancient samples (Fig. 3, Supplementary Fig. 9). Surprisingly, this pangenomic approach also revealed the presence of a hitherto undetected plasmid in the Second Pandemic: pTP33. This rare 34kb plasmid was found to be present in all, but three, of the high coverage Second Pandemic shotgun samples analysed, and is found in all the five subclades with data.

To further investigate how the pTP33 plasmid compares to the three other plague plasmids, we characterised the plasmid copy number for all four plasmids in each sample (Fig. 3c, Supplementary Table 4). This analysis revealed a couple of interesting observations: 1) The two large plasmids, pCD1 and pMT1, occur at low copy numbers, averaging of 3.0 and 1.33 copies per chromosomal genome, respectively in Second Pandemic samples, while the two smaller plasmids, pPCP1 and pTP33, express high copy numbers of 10.0 and 9.5, respectively. 2) preLNBA and LNBAminus plague strains generally have lower plasmid copy numbers than more recent strains. This difference is most striking for pPCP1 where the copy number in LNBA+ strains is more than double that of preLNBA and LNBA-strains (average copy number: 3.8). The increase in pPCP1 copy number occurs at the time of selection for the V51L and I259T *pla* substitutions^21^ and pMT1 copy number increase coincides with the acquisition of the *ymt* gene^22^. 3) Although the average copy number of pTP33 is around 10, there are a handful of outliers with dramatically elevated copy numbers ranging from 17 to 196. All these outliers are from the two most basal and earliest clades in our phylogeny (basal1 and basal2). The two cases of highest copy numbers for pTP33 (196 and 77, respectively), were detected in two samples from the Black Death clade (basal1): CGG100441 from Vor Frue Kirkegård, Denmark and CGG100725 from Tjærby, Denmark.

### The distribution of plasmid PTP33 in space and time

The surprising discovery of pTP33 raises several questions about the distribution of the plasmid in ancient and modern samples. Based on presence/absence data of the plasmid across all published ancient shotgun samples with a coverage over 0.1X (Fig. 4a), we found the plasmid to be present in almost all strains from the Second Pandemic sub-branch, while it is undetectable in all earlier strains from the preLNBA, LNBAminus and LNBAplus clades as well as the Justinianic plague. A similar presence/absence analysis for the modern data revealed that the plasmid was present in 16 out of 2,241 modern plague samples (0.7%) and absent in *Y. pseudotuberculosis* and *Y. similis* (Fig. 4b). Notably, all the pTP33 modern samples fall within that same three small subclades of branches 2-4 (exclusively in subgroups 2.ANT.2, 3.ANT2 and 4.ANT1; Fig. 4b). Geographically, these strains all come from two contained mountainous areas in Central Asia, the Tuva region and Tibet^23^ (Fig. 4e). The modern 2.ANT2 samples from the Tuva region were found in lice, fleas, ticks, long-tailed ground squirrels, Tabagan marmots, and Daurian pika^23^. The louse, flea and tick hosts were not specified. Furthermore, in one case, the plasmid was identified in a sample from a human pneumonic plague case in Tibet, China^23^. As such, the presence of pTP33 lineages across this geographic region, might reflect adaptation to local enzootic transmission cycles, possibly involving flea and/or louse transmission.

**Figure 4.**
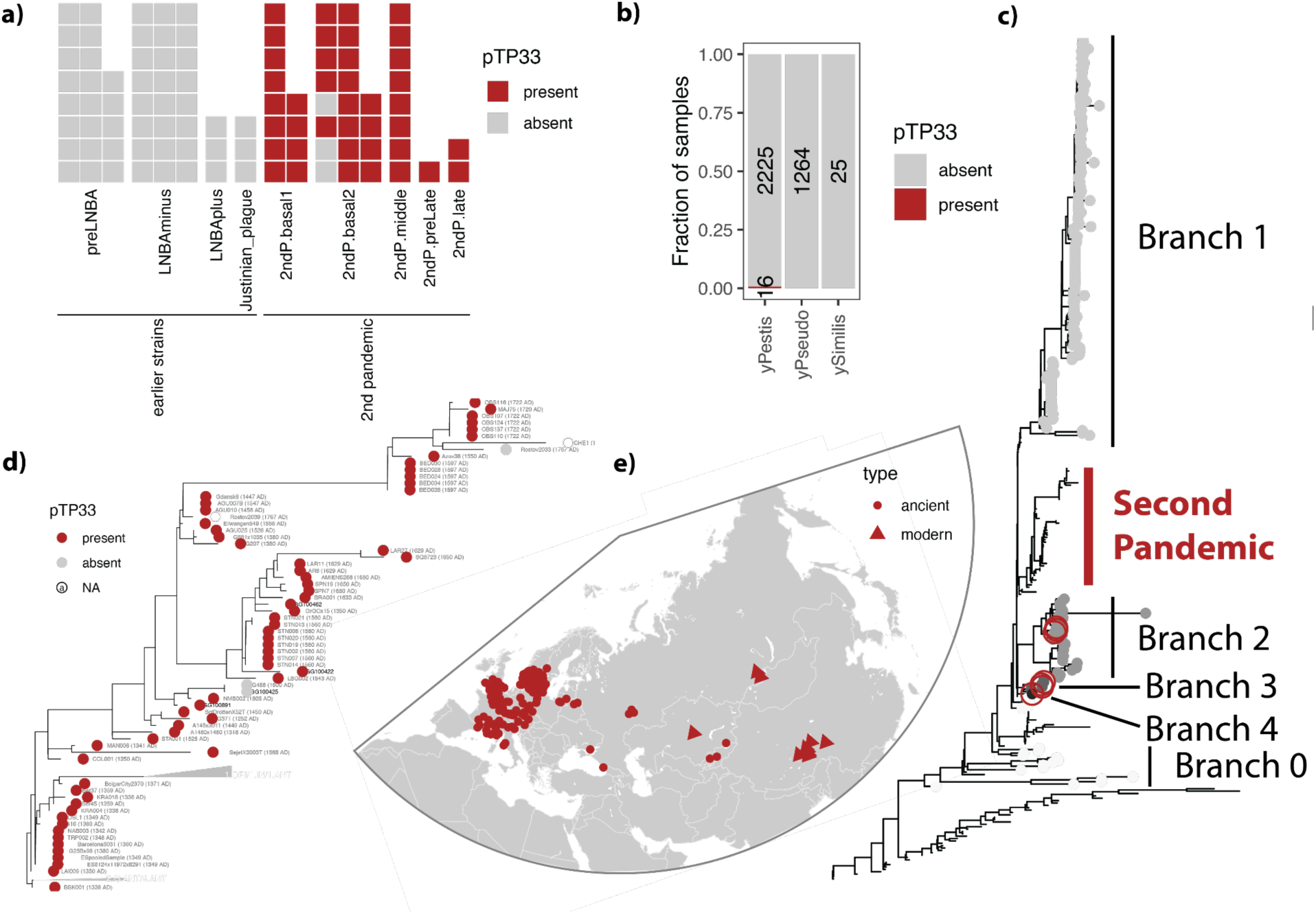
The distribution of plasmid pTP33 through space and time. **a)** Presence/absence of pTP33 in ancient DNA shotgun data with a chromosomal coverage over 0.1X. The plasmid was considered present when the average plasmid copy number surpassed 0.01. **b)** Presence/absence of pTP33 among modern samples within the *Y. pseudotuberculosis* species complex, using the same detection thresholds as for the ancient data. **c)** Distribution of the modern strains carrying the plasmid across the full phylogenetic tree of *Y. pestis*. Modern samples are represented by circles coloured by their branch number, while ancient samples have no tip point shapes. pTP33 positive modern strains are highlighted by red open circles. **d)** Distribution of pTP33 positive samples within the phylogenetic tree of the Second Plague Pandemic using both capture and shotgun data. The detection threshold follows the same logic as described above for shotgun data only (above 0.01 copy number in samples over 0.1X chromosomal coverage), but in cases where only capture data existed for a sample with no evidence of the plasmid, plasmid presence/absence was assigned to unknown (NA). **e)** The distribution of the plasmid across Eurasia in ancient (circles) and modern (triangles) samples.

Having established the overall distribution of the plasmid in shotgun data, we turned to capture data to investigate if trace amounts of off target pTP33 reads might have survived the enrichment washes. While at much lower coverage than on-target sequences, we found that pTP33 is also present at detectable levels in almost all Second Pandemic capture samples (Fig. 4d). Importantly, this analysis demonstrated the presence of pTP33 in all three strains from Kyrgyzstan (BSK001, BSK003 and BSK007) dated to 1338– 1339, representing the earliest known plague cases from the Second Pandemic which are placed in the phylogeny at the node ancestral to the first outbreak of the Black Death^1^. Furthermore, this visualisation identifies the loss of the plasmid twice over the course of the Second Pandemic: once in the 17th century along the ‘2nd.basal2’ branch of G488 (Latvia; 1550-1650 AD), CGG100425 (Denmark; 1600-1800 AD) and the lower coverage sample CGG100404 (Denmark; 1536-1806), and again in the 18th century along the ‘2ndP.late’ branch of Rostov2033 (Russia; 1762-1773 AD) and CHE1 (Russia; 1700-1800 AD). This loss of pTP33 late in the course of the pandemic, follows the same pattern as the depletion of the *pla* gene which occurred at around ∼1400 AD^20^ and has been shown to attenuate disease virulence^13^. Following a similar reasoning, it is possible that the loss of pTP33 might reflect that the transmission route facilitated by the presence of pTP33 was no longer a important for plague in the above regions due to, e.g., distinct local enzootic transmission cycles or improved hygiene or infrastructure.

### Diversity and genetic content of pTP33

Pairwise comparisons of variant positions among all modern and ancient pTP33 sequences indicate very low diversity with a maximum five diverging SNPs detected between two samples (Supplementary Fig. 11). Because of this low diversity, the pTP33 phylogeny generally has low bootstrap support values, suggesting limited phylogenetic structure in the data (Supplementary Fig. 12, Supplementary Fig. 13). To assess if this pattern still holds true when examining genetic variation beyond single nucleotide polymorphisms, we carried out de novo assembly of the plasmid using the data from our highest coverage sample (CGG100891, pTP33 coverage: 98X, Supplementary Fig. 14). With only a single SNP distinguishing the ancient assembly and the five modern pTP33 assemblies, this analysis corroborated our findings of a highly conserved plasmid even at the full assembly level (Supplementary Fig. 15).

Similar to previous reports^24^, we identified 52 open reading frames within the plasmid consisting of 12 ‘hypothetical proteins’ of unknown function, 12 ‘Phage related proteins’ of unknown function and another 28 proteins with known function (Supplementary Table 5). The majority of genes with known function are associated with either maintenance and inheritance (parA, parB, replication initiation protein, and a YefM-YoeB toxin-antitoxin module) and the Phage-derived backbone of the plasmid (integrase/recombinase, terminase, holin, lysozyme, tail/head/baseplate/tail-fiber genes, and other phage-like proteins). However, we also identify exopolysaccharide production protein ExoZ (exodeoxyribonuclease VIII) which has been associated with cell-surface polysaccharide modification and biofilm generation. Interestingly, this proposed link between pTP33 presence and increased biofilm formation has been validated experimentally. In a study comparing the effect of plague plasmid content on biofilm formation, it was demonstrated that the presence of pTP33 was associated with an increased biofilm formation and earlier proventricular blockage in both the rat flea (*Xenopsylla cheopis*) and two other rodent fleas. Unfortunately, these tests were not performed on the human body louse or human fleas^25^.

Taken together, these results support an interpretation of pTP33 as a phage-plasmid-like or plasmidised phage-derived element rather than a conventional plasmid with sporadic phage elements and suggest that the plasmid is involved in cell-surface polysaccharide modification, biofilm formation and faster transmission through proventricular blockage.

### Origin of pTP33

We performed nucleotide-level Blast homology searches to compare each pTP33 coding sequence against the nt database of curated publicly available DNA sequences. We generally found the highest similarity to pTP33 in an *Erwinia* phage LS-2018a/*Erwinia tracheiphila*-associated mobile-element lineage (Supplementary Fig. 16). It is noteworthy that these homologues were found in consecutive loci that encoded functionally related phage proteins such as terminase, capsid, virion/tail and baseplate and lysis-associated functions (Supplementary Fig. 17). This concordant similarity suggests inheritance of a shared mobile-element module because both gene content and local synteny are conserved and suggest a mobile-element sharing or common recent ancestry that contrasts the distant relatedness between *Yersinia* and *Erwinia*. The lower-ranked matches provided information on the broader evolutionary distribution of pTP33-like genes. For example, we identify similarity to *Serratia marcescens* (part of the family Yersiniaceae) and hits to other Enterobacterales such as *Xenorhapdus budapestensis*, *Enterobacter hormaechei,* and *Sodalis* phage phiSG1. Taken together, the pTP33 homologues were concentrated in Enterobacterales with established insect- or invertebrate-associated lifestyle phase, including *Erwinia* (Cucumber beetle gut), *Serratia* (common in several insect guts including mosquitos), *Xenorhapdus* (symbiotic in the gut of Heterorhabditis nematodes) and *Sodalis* (tsetse fly tissues including gut).

To investigate if we could identify pTP33 homologs from metagenomic datasets instead of just relying on the curated sequences of the nt database, we searched public short read archives for reads with high similarity to the coding sequences of pTP33 using Logan^26^ (Methods). Using this approach we identified 48 metagenomes with plasmids similar to pTP33 but with no evidence of *Y. pestis* in the data (Supplementary Table 6). In agreement with the results from the nt database, some of the read archives with the closest similarity to pTP33 comes from the plant pathogen *Erwinia tracheiphila*^27^, and many of the other 48 pTP33 homologs derive from either plant or insect metagenomes suggesting that pTP33 plays a role in bacterial fitness in insect vectors (Supplementary Fig. 18).

These homology searches have important limitations. Sequence similarity alone does not demonstrate evolutionary direction, horizontal transfer history, or biological function and high similarity does not prove that the matched organism was the donor of any specific pTP33 gene. Consequently, for our pTP33 analysis, the most important observation is not that specific genes have particular high similarity hits, but that multiple adjacent genes involved in DNA packaging, capsid formation, virion structure, tail/baseplate assembly, lysis, partition and recombination collectively map to the same *E*. LS-2018a/ *E. tracheiphilia*-associated element and to progressively more divergent homologs in other Enterobacterales. This observation supports the hypothesis that pTP33 is a stably maintained *Y. pestis* phage-plasmid-like element in specific ecological circumstances that is derived from, or share ancestry with, broader phage-plasmid-like elements circulating among Enterobacterales.

That pTP33 is so closely associated with only the Second Plague Pandemic, found in the earliest samples from Kyrgyzstan pre-dating the Black Death, and undetectable in *Y. pestis* from the First and the Third Plague Pandemics is intriguing; it suggests that pTP33 might have been acquired just before the Central Asian diversification of plague into branches 1-4 shortly before the start of the the Second Plague Pandemic and spread together with the Second Pandemic plague lineage. It also suggests that although the third plague pandemic lineage descended from the Second Pandemic lineage, pTP33 was no longer beneficial to plague transmission, possibly because of changes in the vector and animal reservoir that made the plasmid redundant. The persistence in Tuva-Tibet plague lineages could reflect continuous selection within Central Asian wild rodent-ectoparasite systems.

## Discussion

The high mortality of the Second Pandemic in general and the Black Death in particular is indisputable. It is also well established that the Third Pandemic in Europe had a much lower death toll^28^ than its predecessor despite the Third Pandemic plague strain (1.ORI) having evolved from the Second Pandemic branch. This striking epidemiological difference has generally been explained by changes in the human environment rather than by major differences in the plague bacterium itself. For example, it has been proposed that demographic differences between the Middle Ages and the industrial era, such as living conditions, preventive policies, health systems and general hygiene, might explain the disparity between the two pandemics. Our findings, demonstrating the presence of the plasmid pTP33 throughout most of the Second Plague Pandemic and its absence in the Third Pandemic, introduces another possibility: that the transmission ecology of the bacterium itself also differed between the two pandemics.

### Ecology of modern pTP33 carrying strains

Here, we have established (1) that the pTP33 plasmid was likely acquired immediately prior to the outbreak of the Black Death, (2) likely derived from phage-plasmid-like elements common in insect gut-associated bacteria, and (3) was ubiquitous throughout most of the Second Plague Pandemic. We could not detect pTP33 in any plague strain prior to the Second Pandemic, nor in the plague strains associated with the third plague pandemic. Today, pTP33 persists in pockets of modern plague strains of branches 2, 3, and 4 in the Tuva-Tibet plague region of Central Asia. These modern pTP33-carrying strains have been found in a range of infected mammalian hosts and ecto-parasites, including fleas and lice^23^. The Tuva-Tibet plague region encompasses a particularly diverse array of naturally infected animal hosts and specialised ecto-parasite systems compared to, e.g., Madagascar where plague bacteria do not carry pTP33 and plague reservoirs are more geographically and epidemiologically concentrated on a few rodent hosts and a limited number of key flea species, despite a substantial flea fauna^29,30^.

The persistence of pTP33 in the Tuva-Tibet plague strains likely reflects the continued selection for pTP33 within that region’s very diverse wild rodent-ecto-parasite systems. Experimental data have demonstrated that pTP33 appears to increase bacterial biofilm formation and/or aggregation in the gut of the three flea species examined so far, which might increase plague bacteria retention^24,25,31,32^. Consequently, it is possible that pTP33 contributes to vector ecological flexibility rather than to an intrinsically broader animal host range. However, as many flea species can live on more than one animal host, the host range might also be expanded if pTP33 increases the robustness of plague bacterial colonisation, and hence infectious dose, in, e.g., fleas that can live on a range of animals permissive to plague, but without prior host reservoir potential.

We propose that the establishment of these potential new European rodent-flea/louse plague reservoirs might have been critically dependent on pTP33 and hence would not have existed before or after the Second Pandemic. It is noteworthy that the black rat-associated flea, *Xenophylla cheopis*, historically associated with the Second Plague Pandemic, is not rat-specific, but can live on other rodents and small mammals. While *X. cheopis* has been detected in Central Asia, it does not play a major role in the Tuva plague systems. In contrast, two major plague hosts in the Tuva plague region, the long-tailed ground squirrel and the Tarbagan marmot, both carry ectoparasites also found in Europe. The long-tailed ground squirrel often harbors *Linognathoides urocelli* lice, which can live on the European ground squirrel, and Tarbagan marmots carry *Linognathoides palaearctus*, which can thrive on all marmot species from Central Asia to Eastern Europe. Similarly, the flea, *Citellophilus tesquorum*, is found on both marmots and ground squirrels in the Central Asian plague region^33^, and *C. tesquorum* subspecies and related *Citellophilus* fleas extend westwards with ground squirrel hosts into eastern and southeastern Europe^34^. We do not think it is likely that an infected Central Asian host animal was transported to Europe; however, these examples of shared ectoparasites suggest that *Y. pestis* carrying pTP33, could facilitate the spread of plague in diverse European wild and commensal rodent-ectoparasite systems, outside of the of the classical black rat/flea transmission cycle.

### Historical and epidemiological evidence for human ectoparasites as plague vectors

Two epidemiological differences distinguish the Second from the Third Plague Pandemic. First, population mortality was vastly lower during the Third Pandemic, and this difference became greater as settlement size increased. Although Third Pandemic plague could cause high mortality in small communities, mortality remained below 10% in non-evacuated towns of ∼11,000–52,000 inhabitants and declined to ∼2–3% in the largest cities. In contrast, during the Second Pandemic Europe, mortality did not decline as settlement size increased. In Provence in 1720-21, mortality was independent of settlement size across communities ranging from fewer than 100 to 90,000 inhabitants, a pattern also observed in independent Black Death city data. This fits the hypothesis well: if Second-Pandemic transmission was closely tied to humans and their ectoparasites, it could continue to scale as the human population increased, whereas outbreaks that depended more on local rat–flea epizootics would not necessarily do so. (Supplementary Text 3; Supplementary Fig. 20).

Second, plague spread much faster during the Second Pandemic, with contemporary sources describing new outbreaks within days of the arrival of infected individuals or their belongings. For example, in Venice (1575 AD) plague appeared within days in several parts of the city following the arrival of an infected traveller and the dispersal of his clothing and linen^35,36^. In Palermo (1575), new cases appeared within a few days in households receiving people and belongings from infected houses^35^. Lastly, in southern France (1628), plague spread within days of the return of travellers from a plague-affected inn to Fons and Cajarc^37^. These outbreaks are difficult to explain by the classical rat-flea model established during the Third Pandemic, in which human outbreaks follow the die-off of local infected rat populations (rat epizootics). This model predicts a 19-27 day lag from the first plague infection in rats to the first human death and has subsequently been extrapolated to analyses of the Second Pandemic. The rapid establishment of new outbreaks during the Second Pandemic, together with the very different scaling of mortality, is consistent with spread by human ectoparasites, including fleas and lice. (Supplementary Text 4)

These observations are also consistent with systematic analyses of historical texts linking plague to clothing and movable goods, and with modelling of Second Pandemic outbreaks, in which a human-ectoparasite transmission model provided a better fit than rat–flea or pneumonic models in seven outbreaks^38,39^. The Iceland plague epidemic (1494-95 AD) represents a strong case for non-rat associated plague dissemination. Icelandic annals attributed this plague outbreak to cloth brought ashore by ship, an account consistent with transmission by ectoparasites carried in fabric^40^. This interpretation is supported by recent genomic analyses demonstrating the presence of *Y. pestis* in 15th-century Iceland^41^ and the lack of any contemporary historical sources mentioning rats, and no archaeological or ancient environmental DNA evidence for the presence of a rat population in medieval Iceland^42,43^. Together these historical accounts of rapid plague spread and evidence of transmission in the absence of rats suggest a greater role than previously appreciated for human-ectoparasite plague transmission cycles and possibly other animal-ectoparasite cycles. These accounts and results support the idea that pTP33 allowed *Y. pestis* to use a wider range of vectors and/or possibly increased the infectivity of these vectors, including human fleas and lice, which facilitated plague dissemination.

### Possible plague vectors of the Second Pandemic

During the Middle Ages, unstable climatic conditions after ∼1315 together with the population disruptions driven by recurring plague outbreaks caused a reorganisation of farming and trade. These changes might have increased competition and contact between commensal and wild rodents, which in turn increased the likelihood of inter-rodent transmission of infected fleas and lice (see Supplementary Text 2). A plague strain carrying pTP33 might have had a selective advantage if it helped sustain diverse wild and commensal rodent-ectoparasite plague systems in times of disruption of rodent and human communities.

Furthermore, the striking pattern of pTP33 ubiquity and local loss late in the pandemic follows the same course as the depletion of the *pla* gene, which is linked to pneumonic plague and high mortality in both humans and rats^13^. Late Second Pandemic selection for decreased virulence, together with the elevated pTP33 copy number in many Black Death strains, points to a similar central role of pTP33 in the mortality of the disease. However, the role of pTP33 might not be directly related to the disease course in humans and rats but rather linked to increased likelihood of infection and sustained plague spread within Europe. Likewise, the loss of pTP33 in two locations late in the Second Pandemic could suggest that transmission via the black rat/flea cycle in these places was so efficient and specialised that pTP33-linked traits became redundant. Our results thus suggest that the ecology of Y. pestis changes before and during the Second Pandemic as the interplay between the bacteria and host populations were affected by the high plague-associated mortality, which results in changed selection pressures on the bacteria.

Modelling of Second Pandemic plague outbreaks demonstrate that the epidemiology cannot be explained by spread via rats and their fleas alone. Conversely, modelled scenarios of a more direct spread between humans via the human body louse (*Pediculus humanus humanus*) or the human flea (*Pulex irritans*) fit the data more accurately^38,44^. While human fleas can harbour *Y. pestis*, they are not considered an effective vector based on experiments with pTP33 negative *Y. pestis* strains^45,46^. In contrast, human lice have been shown experimentally to be able to carry pTP33 negative plague bacteria in the gut and salivary glands in concentrations sufficient for human infection^47^. Furthermore, the presence of louse-bourne relapsing fewer (*Borrelia recurrentis*) throughout the European archaeological record from the Neolithic to the Middle Ages has robustly established that human body lice were ubiquitous in Europe at the time of the Black Death^48^.

### A proposed mechanism for pTP33

As outlined above there is ample evidence to suggest that the classical plague transmission cycle via black rats and their fleas was less important during the Second Plague Pandemic than the Third. There is also experimental evidence that links pTP33 with increased biofilm formation and earlier proventricular blockage across a diverse selection of vector species. Based on these observations, we hypothesise that the Second pandemic plague strain acquired pTP33 at some point in central Asia before the year 1339 AD, which enhanced bacterial persistence across a broader range of vector species and facilitated transmission among both humans (via body lice and fleas) and a diverse range of animal hosts. Spread of *Y. pestis* carrying pTP33 might furthermore have been especially advantageous during 1346-1700 AD because of the disruption imposed on human and rodent populations due to unstable and colder climate conditions and its consequences for farming, trade and human behaviour. This change in the ecology of the bacteria helps explain the extremely high transmission rates, the prolonged time span of the Second Pandemic relative to the third, and the associated high mortality rates during the Black Death (1346-1353 AD) and the subsequent Second Plague Pandemic (1346-1840 AD). Our findings highlight that even ancient, well-studied bacterial pathogens like plague can undergo transient evolutionary changes through plasmid acquisition that alter pandemic transmission patterns and impact and cautions against uncritically extrapolating epidemiological models across pandemics.

## Methods

### Sampling

Samples were selected to represent medieval and post-medieval populations from different regions of Denmark, with a focus on larger, well-documented burial sites. Selection was further guided by archaeological context, skeletal preservation, and accessibility. Most Danish samples are curated by the Laboratory of Biological Anthropology at the Globe Institute, University of Copenhagen. Permission for destructive sampling was obtained from the museums owing the respective collections.

### Drilling, Library building, and sequencing

All samples were processed in the ancient DNA clean laboratory facilities at Globe Institute, University of Copenhagen following strict ancient DNA guidelines^49^. Tooth samples were subsampled by removing the exposed surface of the root using a metal drill disc before cutting off the root. Roots were then split in half vertically, and the pulp cavity and dentine was drilled out to enrich the sample for cementum^50^. For petrous bones, the inner otic capsule was isolated using a small drill disc, drilling away all porous bone material surrounding the otic capsule. For both sample types, bone material was then crushed with a pair of pliers and split into two subsamples aiming for around 80-160 mg of material in each.

DNA extractions were carried out using the automated laboratory processing pipeline available at Globe Institute, University of Copenhagen following the protocol described in ref.^51^, with two initial demineralisation steps described in ref.^52^. Following extraction, the majority of DNA extracts were split in two to create two batches, one subjected to USER enzyme (New England Biolabs) treatment and one that was left untreated (Supplementary Table 7). Library building was then carried out using either the double stranded library build approach described in ref^53^ or the Santa Cruz Reaction single stranded library build protocol^54^. Both protocols were adapted for the Biomek i5 Automated workstation (Beckman Coulter)^52^. Lastly, following index PCR, libraries were sequenced on the NovaSeq 6000 instrument (Illumina).

### Capture enrichment

High complexity sequencing libraries that had not reached saturation of plague DNA yet, were subjected to in-solution hybridisation capture enrichment using previously published^55^ myBaits probes (Daicel Arbor Biosciences, Supplementary Table 7). Enrichment was carried out following the manufacturers High Sensitivity protocol in pools of approximately 8 libraries using a single round of capture. For each pool, equimolar amounts of each library were pooled aiming for 500ng of total input DNA. Post-capture reamplification was carried out in 50 μL reactions for 16 cycles using IS6 (P7) and IS5 (P5) primers and the Kapa U+ master mix (Roche).

### Data preprocessing and pathogen screening

All sequenced data was preprocessed using AdapterRemoval (v.2.3.2)^56^ set to remove adapters, trim ambiguous bases at read ends and to collapse paired reads overlapping with more than 10bp. To characterise the microbial diversity within each sample, we screened all preprocessed reads using the ‘pathopipe’ tool (https://github.com/martinsikora/pathopipe)^48^ on the krakenUniq database ‘hum_microbe_release_20250428’ (available from: [link](https://erda.ku.dk/archives/1d29e091b69cabe43093440eeb396212/diseases/public_supplementary_data_repo/hum_microbe_release_20250428.tar.gz)). We identified plague samples by parsing the summary files from the pathogen screening, classifying a sample as plague positive when the following criteria were met for *Y. pestis*: number of reads over 50 (nReads>50), best species hit within the genus (krakenKmerRank==1), an observed coverage versus estimated coverage ratio over 50% (coveragePRatioCorr > 0.5), an average nucleotide identity over 97% (ani>0.97), and an average softclipping of less than 8bp (nSoftClipAvg<8).

### Linear mapping to CO92

Preprocessed data of all plague positive samples (n=110) was mapped to a reference genome consisting of the CO92 assembly (GCF_000009065.1) with the pTP33 sequence (NZ_CP178016.1) from the GCF_046895395.1 assembly appended. Mapping was carried out using bowtie2 (v.2.3.2)^57^ with the following mapping parameters: -D 20 -R 3 -N 1 -L 20 -i S,1,0.50 --end-to-end --no-unal. The resulting alignment files were merged to library level and sorted with Samtools^58^ (v.1.12), followed by flagging of PCR- and optical duplicates using MarkDuplicates (v.2.27.4) from Picard tools. Lastly, alignments were merged to sample level, followed by the removal of previously flagged duplicates and alignments of low quality (mapping quality < 30). Using thse filtered alignment files, we calculated per gene average depth of coverage using mosdepth for all known virulence across all ancient samples and compared these to representative modern samples of the major plague clade found in the modern form of the disease (Supplementary Fig. 19).

Variants were called using the joint genotyping approach from GATKs (v4.3.0.0)^59^. First, pr-sample genomic variant call format (GVCF) files were generated from the filtered alignment files using HaplotypeCaller set to evaluate all positions of the reference genome (ERC BP_RESOLUTION). The resulting GVCF files were merged using CombineGVCFs, followed by joint genotype calling with GenotypeGVCFs. Next, using GATKs VariantFiltration genotypes not meeting the following thresholds were filtered out by setting the call to N: Low depth (DP<3), high depth (DP>1000), low genotype quality (GQ<50) or poor genotype support (AB<0.9). Lastly, we filtered out poor quality variants not meeting the following criteria: QD < 2.0, FS > 60.0, MQ < 40.0, MQRankSum < -12.5, ReadPosRankSum < -8.0.

### Phylogenetic analyses

In order to reconstruct the plague phylogeny, we restricted the variant call file described above to SNPs using GATKs SelectVariants, and applied these variants to the chromosome of CO92 (NC_003143.1) in a sample-wise manner using bcftools consensus (v1.16)^60^. We used the resulting multi-fasta to reconstruct the plague phylogeny using RAxML-NG (v. 1.2.0) with a GTR+G substitution model and the *Y. pseudotuberculosis* genome from serotype O:1c (SAMEA7160327) as the outgroup. We also calculated bootstrap support values for the predicted phylogeny using RAxML-NGs two built in bootstrapping metrics based on the transfer bootstrap expectation and Felsenstein’s bootstrap proportions. Lastly, we visualised the best phylogenetic tree using the ggtree (v.2.0.4)^61,62^ package in R (v.4.2.2).

For lower coverage samples (0.01X - 4X), we characterised phylogenetic associations using the placement algorithm implemented in EPA-ng (v0.3.8)^63^. Briefly, we created a SNP only fasta file, by subsetting the multi-fasta file described above to positions of all SNPs in our variant call file using bedtools getfasta. We ran EPA-ng on the resulting fasta file using the best tree and model from RAxML-NG described above including a maximum of 1000 placements per sequence in the final output (--filter-max 1000). Lastly, we visualised the resulting placements on the phylogenetic tree using ggtree (v.2.0.4)^61,62^, and assigned each sample to phylogenetic groups based on a visual inspection of the resulting plots.

### Pangenomic characterisations

The input data for pangenome variation graph building was generated by downloading all complete genomes of the taxa *Y. pestis*, *Y. pseudotuberculosis* and *Y. similis* using NCBI’s command line tool ‘datasets download’. Plasmid sequences were then selected by extracting all contigs with ‘plasmid’ in the fasta header. Next, the pangenome variation graph was built using these complete plasmid assemblies as input for pggb^64^ with default settings except from the use of a segment length of 500bp. In order to classify all plasmids in the graph despite differences in naming schemes used in different research fields, host animals and bacterial species, we carried out sequence partitioning on the resulting graph using the two python scripts paf2net.py and net2communities.py from pggb. Using this approach we detected 8 distinct communities in the graph, representing the following plasmids: pMT1 (225 sequences), pCD1 (240 sequences), pCRY (9 sequences), pPCP1 (204 sequences), pTP33 (7 sequences, including the pS96127 plasmid identical to pTP33^23^), pCKF (3 sequences), pVM82 (2 sequences) and pYPSE1-3 (2 sequences). Moreover, 6 additional plasmids were characterised as singletons and were not part of any community: Unnamed_1 (NZ_CP155630.1), P_59kb (NC_009704.1, pYpstbSP1303.1 (NZ_CP130904.1), pYpstbSP1303.2, (NZ_CP130905.1), pYPSE1-1 (NZ_AP028969.1), Unnamed_2 (NZ_CP007231.1).

Graph mapping was carried out using vg giraffe^65^ on the plasmid graph indexed with vg autoindex. After mapping, gam files were merged at the library level retaining only alignments with mapping quality over 1, followed by duplicate removal using the custom made python gaf_rmdup.py. Lastly, gam files were merged at sample level, and basic coverage statistics for all nodes in the graph were calculated for alignments with mapping quality over 30 and a normalized alignment score of at least 0.9.

We characterised each node in the graph based on the sequences traversing that node. Nodes traversed exclusively by sequences of a single plasmid community were assigned to that community, whereas nodes found across different plasmid communities were characterised as ‘shared diversity’. Within each group, we defined a core genome represented by all nodes present in more than 33% of the sequences within that group. For each ancient sample, we characterised a node as being present when the average coverage across that node was over half the average coverage for that sample. We visualised the coverage of ancient samples by counting the length of nodes with coverage out of the total length of the core genome for each community (see Fig. 3b).

To calculate the copy number for each plasmid in each ancient sample, we calculated average coverage of the core genome of each plasmid community for each sample. Next, we normalised this coverage to the average coverage of the chromosome, as calculated from the linear mapping to CO92 described above.

### Logan-Search

We applied the k-mer search engine Logan Search^26^ on the coding sequences of NZ_CP178016.1 to identify public short read archives with sequences of high similarity to pTP33. We split the NZ_CP178016.1 sequence into coding sequences using bedtools getfasta^66^ (v2.31.0), on RefSeq genes specified in <u>GCF_046895395.1_ASM4689539v1_genomic.gff.gz</u>. We submitted each gene sequence to Logan Search individually, searching against the ‘all’ database and extracted all biosample-ids from the results table upon completion of the job. Next, we downloaded fastq data for all identified biosamples with similarity to pTP33 using enaDataGet (1.7.1), and preprocessed the data with fastp^67^ (0.23.4). We mapped the resulting reads to our reference genome consisting of the CO92 assembly with the pTP33 sequence appended as outlined above. Based on the depth of coverage (cutoff: 1X) and the coverage evenness statistics (cutoff: 0.3) from the mapping results we characterised each biosample as either positive for plague and pTP33, positive for pTP33 with no evidence of plague, or none of the above. To visualise the diversity of all biosamples were pTP33 was detected, we generated a pTP33 phylogeny using raxml following the approach described above and visualised the results with ggtree.

### De novo assembly

We assembled the metagenome from our highest coverage plague sample (CGG100891) using MEGAHIT (v1.2.9)^68^ with default settings on all human depleted data from CGG100891. The resulting unfiltered assembly consisted of 155,6245 contigs with an N50 of 514 spanning a total of 891,085,464 base pairs, which was filtered to keep contigs over 3kb and converted into a reference genome. Next, we mapped all reads from CGG100891 back to the filtered assembly and calculated basic mapping statistics, including average depth of coverage, breadth of coverage and depth of coverage across the sequence for each contig. To identify contigs deriving from plague, we extracted reads from CGG100891 that mapped to the chromosome, pCD1, pPCP1, pMT1 or pTP33, respectively, in the linear mapping, and mapped these to all assembled contigs over 3kb. We classified contigs as deriving from one of these genetic elements if any reads from that element mapped to the contig in question. Using this approach, some contigs were classified as deriving from multiple plasmids because of repetitive elements present across different elements. In these cases, we assessed the coverage across the contig manually (Supplementary Fig. 14), and assigned the contig to the genetic compartment that most of its sequence was derived from.

## Supplementary Information

### Supplementary Text

#### Supplementary Text 1 - site descriptions

##### St Clemens and St Nicholas, Sweden

During the early Middle Ages, the town Helsingborg had three parish churches, all situated on the coastal ridge (*landborgen*) one of them was St. Clement church. It was in use from approximately 1050 to 1536. Analyses of the burial remains from St. Clement’s revealed a marked decline in the number of interments during the High and Late Middle Ages. This is attributed to the relocation of the town center and a construction of the establishment of the Dominican friary dedicated to St. Nicholas shortly before 1275.

St. Clemens cemetery holds a cross-section of the city’s population, whereas the graves at the Dominican friary were predominantly purchased by the wealthy. This is reflected in differences in stature: those buried at the friary were significantly taller than those at St. Clemens. St. Clemens is also the site where the cases of leprosy in the town were found-a disease associated with poverty, while the Dominican friary yields cases of gout, a condition associated with affluence

##### Niels Ebbesens Vej 10 (KBM4406)

The site at Niels Ebbesens Vej 10 (Skolen ved Søerne) was archaeologically excavated, revealing at least nine burials. One burial contained the remains of a fetus, estimated to have been approximately 4–5 months of gestational age. The burials were located within a single large burial pit, with coffins arranged approximately three abreast and in at least three superimposed layers. The eastern and western limits of the pit were identified during excavation, whereas its full extent to the north and south remains unknown.

Parts of the burial pit had been disturbed or destroyed during construction of the school in the late nineteenth century. Material originating from these disturbed graves was likely deposited in the remaining sections of a former canal immediately adjacent to the burial site. Based on the archaeological context and contemporary written records, the burials are attributed to the Copenhagen plague outbreak of 1711.

##### Åbenrå 16 (KBM4099)

The Museum of Copenhagen conducted an archaeological excavation at Åbenrå 16 in 2015. A section of an assistant cemetery for Trinitatis Church was uncovered. The cemetery was established in connection with the 1711 plague in Copenhagen and remained in use until 1760. A total of 11 coffins and skeletons from at least 19 individuals were recovered. The coffins were poorly preserved, but the skeletons were all well-preserved. There was a nearly equal distribution of children and adults. Signs of biological stress were observed in several individuals, which may be the result of malnutrition.

The cemetery was located in the southernmost part of the investigated area, and several of the coffins were dug into layers related to the 13th-century medieval rampart, which ran through the plot. A section of the medieval rampart was uncovered together with a remnant of the former street that ran along the inner side of the rampart.

##### Skt. Clemens Kirkegård (KBM3621)

The excavation was conducted on the northern part of the churchyard belonging to one of the eldest known churches of Copenhagen. The church of St. Clemens is first mentioned in the written sources in 1192, but burials at the cemetery have proven older.

1,048 burials were excavated, a small proportion of which contained more than one individual. A few of the deceased were interred in masonry-lined graves, while the majority were buried in pits sized to fit the individual and in wooden coffins.

Both adults and children were found in the cemetery, with a predominance of women. The excavated section of the cemetery was located north of the St. Clemens Church and represents only a small part of the entire burial ground. The burials date from the 11^th^ century until the mid-1500s. The long timespan was shown in the burial intensity of the area where burials often were cut by later multiple grave cuts.

##### Klostergade and Rådhustorvet, Vejle (VKH672, VKH1586, VKH8029)

In 2019–2020, Vejlemuseerne excavated 100 individuals from the Dominican friary cemetery; excavations in the 1980s and 1990s had revealed a further 87. The skeletons were well preserved and predominantly male, with females comprising only one third. The estimated mean age at death among adults was approximately 38 years. The cemetery adjoined the friary, most probably founded around AD 1330–1340 according to written sources. Archaeological evidence provides some support for this chronology: a boundary ditch stratigraphically predating the graves was dated by AMS radiocarbon analysis to the 13th century. The friary closed in AD 1529, when King Frederick I took it over shortly before the Protestant Reformation. In AD 1531, the Crown donated the property to the town of Vejle, and the cemetery was converted into the town square, streets and gardens. Its period of use is thus well defined, extending from approximately AD 1330 to 1531.

##### Kirkegade, Vejle (VKH7816, VKH7864)

Excavations at the cemetery of Skt. Nicolai Church in 2017 revealed 51 well preserved individuals dating to the medieval or early modern period. The sex distribution among the assessed adults was more balanced than at the Dominican friary cemetery. The estimated mean age at death among adults was approximately 39 years. The cemetery is broadly dated from the High Middle Ages to the early 19th century. Construction of Skt. Nicolai Church began in AD 1250 and is believed to have been completed by AD 1256. The earliest building remains excavated in medieval Vejle, just south of the church, may date to as early as approximately AD 1100–1200. The cemetery closed in 1825, and the more recent burials were relocated. Thus, the cemetery at Skt. Nicolai has a broader chronological range and a longer period of use than the Dominican friary cemetery.

##### Holmens Kirke, Denmark (NM538-2013)

In 2013, the National Museum of Denmark excavated a 140 m² area south of Holmens Church in Copenhagen, ahead of the construction of a new parish building. The excavation investigated part of the churchyard, which was in use from 1649 until around 1800.

The earliest activity dates to around 1550, when the area consisted of shallow coastal waters. A low embankment of turf and sand was constructed as part of land reclamation associated with the establishment of a naval shipyard and anchor forge. The forge was built in 1562 and was later converted into Holmens Church. In 1649, the surrounding area was established as a cemetery, which remained in use until around 1800.

A total of 172 graves were excavated. Anthropological analysis identified 166 individuals from defined burials: 147 adults and 19 subadults. Of the adults, 56 were male, 72 female and 19 of undetermined sex, and many had reached an advanced age. The subadults comprised five infants, ten children and four adolescents. Disarticulated remains represented a further minimum of 353 individuals, reflecting the intensive use of the cemetery.

##### Tjærby deserted church and cemetery (KHM 0899), Denmark

Rural cemetery ca. 5 km east of Randers on the north side of Randers fjord, excavated by Kulturhistorisk Museum Randers from 1998 to 2010. The excavation area revealed a stone church and a cemetery containing ca. 1200 graves dating from ca. 1050-1536 AD. The cemetery area measured ca. 45×42 meters (1900m2) and has been fully excavated. In nearly half of the graves, no skeletons were preserved. The remaining burials (673 in total) contained skeletons of varying degree of preservation due to the varying soil conditions in the area. Preservation ranged from few poorly preserved fragments of bones to well preserved and nearly complete skeletons. The best preserved skeletons were found south of the church. Evidence of wooden coffins were found in some of the graves, but the majority of the individuals had been buried without. 57% of the graves with preserved bones contained adults (male and female), 10% subadults and 33% children.

Hyldgaard, I.M, Jørkov, M.L. and Klassen Lutz. *Tjærby Ødekirke og Kirkegaard*. Museum Østjyllands Publikationer vol. 3 (Aarhus University Press, 2016)^69^

##### Chevilly-Larue, France

The necropolis of the Church of Sainte-Colombe in Chevilly-Larue contains the largest collection of rural skeletal remains in the Val-de-Marne department. Archaeological work carried out by the department between 1981 and 1998 revealed 306 primary inhumation graves and the remains of 643 individuals. Carbon-14 dating performed on approximately ten skeletons spans the 9th to 17th centuries, with a predominance of dates falling between 887 and 1030 AD and between 1295 and 1419 AD. Analysis of the osteoarticular and alveolodental pathocenosis reveals infectious, degenerative, metabolic, and neurological diseases, as well as significant vertebral deformities and malformations. Dental lesions of either localized (calculus, caries, periodontal disease) or systemic origin (rickets, congenital syphilis) were also documented. Regarding craniofacial architecture, the prevalence of brachycrany (27%) is comparable to that observed in the population of La Queue-en-Brie (Hadjouis 2018; Hadjouis et al. 2015). Notable vertebral deformities and malformations include spondyloarthropathy, ankylosing spondylitis, spondylolisthesis, and Scheuermann’s disease (Hadjouis 1999, 2026).

Hadjouis, D. 1999. Les populations médiévales du Val-de-Marne. Dysharmonies cranio-faciales, maladies bucco-dentaires et anomalies du développement dentaire au cours du Moyen Age. Coll. Paléoanthropologie et Paléopathologie osseuse, n·1, Eds Artcom’ Paris, 172 p.

Hadjouis, D. 2018. Atlas et traumatismes du monde médiéval et moderne. Préface Yves Coppens, ISTE éditions Londres, 278 p.

Hadjouis, D., Huard, P., Lichon, A-A., Pittion, C. 2015. La nécropole de l’église Sainte-Colombe de Chevilly-Larue (Val-de-Marne) : Population et état sanitaire du Haut et Bas Moyen Age. Actes du Colloque Dynamique des peuplements, modes d’habitat et influences culturelles dans le Sud Est de Paris du Néolithique ancien à la période moderne, Créteil 3-4 juin 2010, supplément n° 3 à la Revue Archéologique d’Île-de-France

Hadjouis, D. 2026. Le rachis humain, du passé au présent : Posture, déformations, malformations, Société Bioprogressive de Ricketts, n° 19, pp. 47-55.

##### La Queue-en-Brie

The village of La Queue-en-Brie is located approximately 20 km southeast of Paris on the Brie plateau, at an elevation of 100 meters. The town was fortified in 1269, but between 1430 and 1652, the village and its church suffered fires and massacres—first at the hands of the English, and later by Huguenots aided by German mercenaries in 1563. The Saint-Nicolas parish was established in 1087. Excavations have unearthed 315 skeletons in situ. The bodies were primarily buried in wooden coffins or simply wrapped in shrouds; in some cases, they were placed in pits roughly hewn into the limestone. Carbon-14 analysis indicates that the burials span the period from the 11th to the 17th century (specifically between 1000 and 1276 AD). The study population comprises 112 women and 81 men, with a very low proportion of juveniles (13%). In terms of craniofacial morphogenesis and architecture, this population differs from other medieval populations in two key respects: greater maximum cranial width and significant basicranial flexion during growth. These features suggest underlying causes rooted in a hereditary and familial combination of specific growth patterns. Indeed, the prevalence of craniostenosis (scaphocephaly, brachycephaly, trigonocephaly, and plagiocephaly) and significant traumatic and congenital hip dislocation points to endogamy (Hadjouis, 2000, 2006; Cho and Hadjouis, 2005).

Hadjouis, D. 2000. Etiopathogénies des dysharmonies cranio-faciales de populations médiévales du Val-de-Marne. In. Andrieux, P., D. Hadjouis, A. Dambricourt-Malassé, Actes du Colloque de Créteil, pp. 170-186.

Hadjouis, D. 2006. Variations anatomiques posturales chez les populations franciliennes du Moyen-Age (Val-de-Marne). L’exemple des dysplasies luxantes de la hanche et des malformations articulaires du genou. Biom. Hum et Anthrop., Paris, 24, 3-4.

Cho, K-H et Hadjouis, D. 2005. Les asymétries cranio-faciales de la nécropole médiévale de la Queue-En-Brie (Ile-de-France, Val-de-Marne). Biom. Hum et Anthrop., Paris, 23, 1-2, p. 95-104

#### Supplementary Text 2 - Historical perspectives

##### Ecological instability and disruptions before and during the Second Pandemic

Historically, ecological conditions in Europe were comparatively more stable before and after the Second Plague Pandemic than those prevailing during the pandemic. High medieval Europe (∼1250 AD) was characterised by high population densities and extensive farming. Commensal black rat populations were well established and coexisting with wild rodents. Around 1315-22 AD, the climate became colder and wetter resulting in major disruptions in the agricultural and rodent food systems. After a short climatic recovery period, cold summers again prevailed ∼1345-1347 AD, resulting in food shortages and long-distance movement of grains and other commodities. The Black Death (1346-1351 AD) resulted in high mortality, collapse and/or reorganisation of farming and sometimes abandonment of settlements and fields^4,70^. This is likely to also have affected the interactions between wild and commensal rodent populations, disrupting old and forming new food networks as less human-associated food was available to commensal rodents. Such disruption might have increased contact and conflict between wild and commensal rodents and consequently increased the likelihood of inter-rodent transmission of infected fleas and lice. Subsequently, the ‘Little Ice Age’ (∼1400-1700 AD)^71^ followed with changed land use and repeated outbreaks of plague. Brown rats expanded in Europe ∼1700 AD and often outcompeted black rats. A plague strain carrying pTP33 might have had a selective advantage if it helped sustain diverse wild and commensal rodent-ectoparasite plague systems in times of disruption of rodent and human communities.

#### Supplementary Text 3 – Settlement-size scaling of plague mortality during the Second and Third Plague Pandemics

We asked whether plague killed a smaller share of the population in larger settlements. For the Second Pandemic, we used Biraben’s data from the Provence epidemic of 1720–21^72^. The dataset contains 72 affected localities, ranging from villages with fewer than 100 inhabitants to Marseille with about 90,000. Mortality was calculated from the reported population and plague deaths. There was no relationship between settlement size and mortality. A tenfold increase in population changed mortality by only +0.001 percentage points (P = 0.9998; Supplementary Fig. 20). Biraben reached the same conclusion from the historical material, stating that there was no apparent correlation between the size of a locality and the severity of the epidemic^72^.

For the Third Pandemic, we used Condon’s data from 13 settlements in western India in 1897– 98^73^. Here the pattern was different. Mortality fell as settlement size increased. A tenfold increase in population was associated with a decline of 7.35 percentage points in mortality (95% CI −11.93 to −2.77; P = 0.0047). This result was not caused by the unusually high mortality in Karve, where 41.4% of a population of 851 died. Removing Karve made the relationship stronger rather than weaker: mortality then declined by 5.47 percentage points per tenfold increase in population (P = 7.5 × 10⁻⁵). In leave-one-out analyses, the relationship remained negative and significant regardless of which Third-Pandemic settlement was removed. The pattern also persisted when both datasets were restricted to the population range they shared. Provence was still flat (+0.42 percentage points per tenfold increase, P = 0.93), whereas mortality in the Third-Pandemic settlements still declined with settlement size (−8.96 percentage points, P = 0.048).

Independent datasets show the same broad contrast. In the Black Death city data of Jedwab, Johnson and Koyama, mortality was likewise unrelated to city size^74^. During the Third Pandemic, Greenwood independently found that infected small villages in Punjab generally experienced higher proportional mortality than larger villages^75^. The comparison concerns how population mortality changed with settlement size, rather than absolute mortality levels between the two pandemics. The Provence data describe completed local epidemics, whereas Condon’s figures cover outbreaks during 1897–98 within a pandemic that continued in India. Together, the data show a simple contrast: Second-Pandemic mortality was independent of settlement size, whereas Third-Pandemic mortality declined as settlement size increased.

#### Supplementary Text 3 – Rapid spread of plague between households and the latency of local rat epizootics

According to Benedictow’s rat–flea model, when infected rat fleas are brought to a new area they first infect the local rats and thus start an epizootic. He gives an estimated time span of 16 to 23 days between the introduction of plague into a rat population and the first human cases, and 19 to 27 days between introduction and the first human death (Benedictow, 2024, pp. 48, 162)^2^. Benedictow also suggests, in the case of the Second Pandemic, that the rat fleas, having been carried in clothing or personal belongings, first infect the rats at the destination before any human cases occur (Benedictow, 2024, p. 449–450)^2^.

The Indian Plague Research Commission found that rat fleas could be carried in clothing and bedding from houses affected by plague and still remain infective; it also stated that imported rat fleas would tend to bite local rats and could thus cause a new epizootic (Advisory Committee Appointed by the Secretary of State for India, the Royal Society, and the Lister Institute, 1907, p. 889–890, 894)^76^.

The outbreaks in Venice and Chiuso during the second plague pandemic allow direct comparison with these intervals. The remaining cases show whether the same pattern of plague following people, clothing and household goods into new households or localities recurs elsewhere.

##### Venice, 1575

Cornelio Morello records that a traveller from plague-affected Val Sugana arrived in Venice on 25 June 1575 and lodged in San Marziale. He died on 2 July. Within a few days, three women in the household had died and two others were ill. Most of his clothing had meanwhile been sold or pawned, and members of the household continued to mix with others. Morello then reports that, again within a few days, plague was discovered in various parts of the city^77^. Palmer adds that the traveller was moved after falling ill, some of his linen reached the convent of Santa Sepolcro, and three foci had appeared in separate districts by the time the Health Office recognized the danger. Their locations corresponded with the movements of the traveller and his belongings^78^ (Palmer, 1978, p. 211–213). Morello does not provide individual dates for each focus but states that the dispersal of clothing, contacts, and the outbreak of plague at several new locations took place over a few days rather than the several weeks needed for a full new rat epizootic to develop at each site.

##### Palermo, 1575

According to Giovanni Filippo Ingrassia, people avoided quarantine by concealing illness in the household, taking their belongings with them and sending members of the family to relatives and friends. He states that afterwards the plague soon appeared “in one house and then in another” at all the places to which they had gone (Ingrassia, 1576, p. 69–70). He doesn’t state whether the first cases in the receiving houses were newcomers or existing residents and therefore the passage provides no reliable secondary-transmission interval. It does note that plague appeared within a few days in several households reached by people and possessions from an infected house.

Ingrassia also notes that of the eighteen purificatori who handled goods from plague houses, seventeen had been admitted to the plague hospital within a few days; in some cases, they had even worn clothes that belonged to plague victims (Ingrassia, 1576).

##### Figeac, Fons and Cajarc, 1627–28

Niall Ó Glacáin attributed the outbreak at Figeac to a muleteer from Auvergne who lodged at the inn of a man named Cleirac and died there. Residents of Fons and Cajarc who had stayed at the same inn returned home and, according to Ó Glacáin, unknowingly infected their towns^79^ (O’Glacan, 1629, p. 12–13). He also says that neighbouring places were contaminated by Auvergnat beggars.

##### Chiuso, 1629

Alessandro Tadino provides the shortest explicit time interval. Although Chiuso had only about forty hearths it housed some 3,000 German soldiers, a number of whom were already ill. Eight of the soldiers who lived in the house of Melchione Castagna died with buboes and other symptoms of the plague. Deaths then took place in the house of Pasino; the next day Pasino’s wife and three of his children died. According to Tadino, within eight days all the relatives and friends who had visited or looked after that household had died, along with some of the members of Francesco Lantera’s household who had been to the affected houses. He characterises the disease as spreading from family to family^80^ (Tadino, 1648, p. 32–33). In this case death occurs within less than eight days, as opposed to Benedictow’s figure of 19 to 27 days between the introduction of the disease into a new rat population and the first human death. Since Chiuso was a very crowded place and more than a hundred soldiers could be lodged in a single house, it is possible that different households shared the same source of infection. Nevertheless, Tadino includes the deaths of visitors, carers and members of another household within that eight-day period.

##### Abbazia di Mandello, 1629

Near Mandello, a family stripped clothing from a dead German soldier. The wife washed the garments and died first, followed by her son, husband and daughter. The daughter had meanwhile stayed in the house of Evangelista Mazza. Tadino says she infected the entire household, where five people died, together with the gravedigger; eight houses in the locality were eventually affected^80^ (Tadino, 1648, p. 43). The sequence goes from the clothing of a dead soldier into one household and then from a member of that household to another; Tadino does not give an interval between the daughter’s stay at the Mazza house and the subsequent cases.

##### Giuseppe Bonfante, Milan, 1629

Giuseppe Bonfante travelled from Cernusco Monte di Brianza to the inn of SS Gervasio e Protasio in Milan and died suddenly after three days. His death was concealed, his body stripped, and his clothes divided between the innkeeper, the Antiano and the gravedigger. Plague later appeared among people associated with the inn (Tadino, 1648, p. 53–54)^80^. The three-day interval ends with Bonfante’s death; Tadino gives no interval to the later cases.

##### Nijmegen, 1635–37, and Amsterdam, 1663

Isbrand de Diemerbroeck repeatedly describes plague travelling with people, clothing and household goods. He writes that experience had often shown travellers from infected places carrying contagion in their clothes or belongings to other cities and regions. He also linked the Amsterdam outbreak of 1663 to travellers arriving from Algiers and Smyrna who first infected their lodgings, from which plague spread further through the city (Diemerbroeck, 1665, Book II, ch. III)^81^. At Nijmegen, goods from plague houses were often sold on the day of a burial, the following day, or within a few days. Diemerbroeck describes a woman who bought a bed-cover, placed it on her bed, and was, together with her husband, infected that first night. In another case, a soldier bought a garment for his son; the boy developed plague, followed by the father, mother and remaining children, all of whom died (Diemerbroeck, 1665, Book II, ch. III)^81^. He also says that he had on many occasions seen straw on which plague patients or the dead had lain capable of transmitting the disease to other people, and mentions clothing, linen and household furnishings as usual carriers of contagion between houses, towns and regions (Diemerbroeck, 1665, Book II, ch. III)^81^.

##### Interpretation

Venice and Chiuso offer the most clear-cut evidence of the timeline. In Venice, plague appeared at a number of new foci within a few days of the dispersal of an infected traveller’s belongings and subsequent contacts involving members of his household. In Chiuso, the relatives and friends who had visited or looked after the Pasino family, along with some members of another household, died within less than eight days.

The same movement-associated pattern is found at Palermo, Figeac, Fons, Cajarc, Mandello, Milan, Nijmegen and Amsterdam; in these cases, plague follows people, their clothing, bedding or household goods into new houses or areas. The Venice and Chiuso sequences are too fast for a complete new local rat epizootic to have preceded human plague at every successive focus.

### Supplementary Table Legends

**Supplementary Table 1**. **Sample metadata and summary statistics.**

**Supplementary Table 2. Overview of published plague samples.**

**Supplementary Table 3. Public *Yersinia* plasmid assemblies used for variation graph building.**

**Supplementary Table 4. Plague mapping statistics.**

**Supplementary Table 5. Bakta gene annotations**

**Supplementary Table 6. Logan results**

**Supplementary Table 7. Sequencing library information**

### Supplementary Figures

**Supplementary Figure 1.**
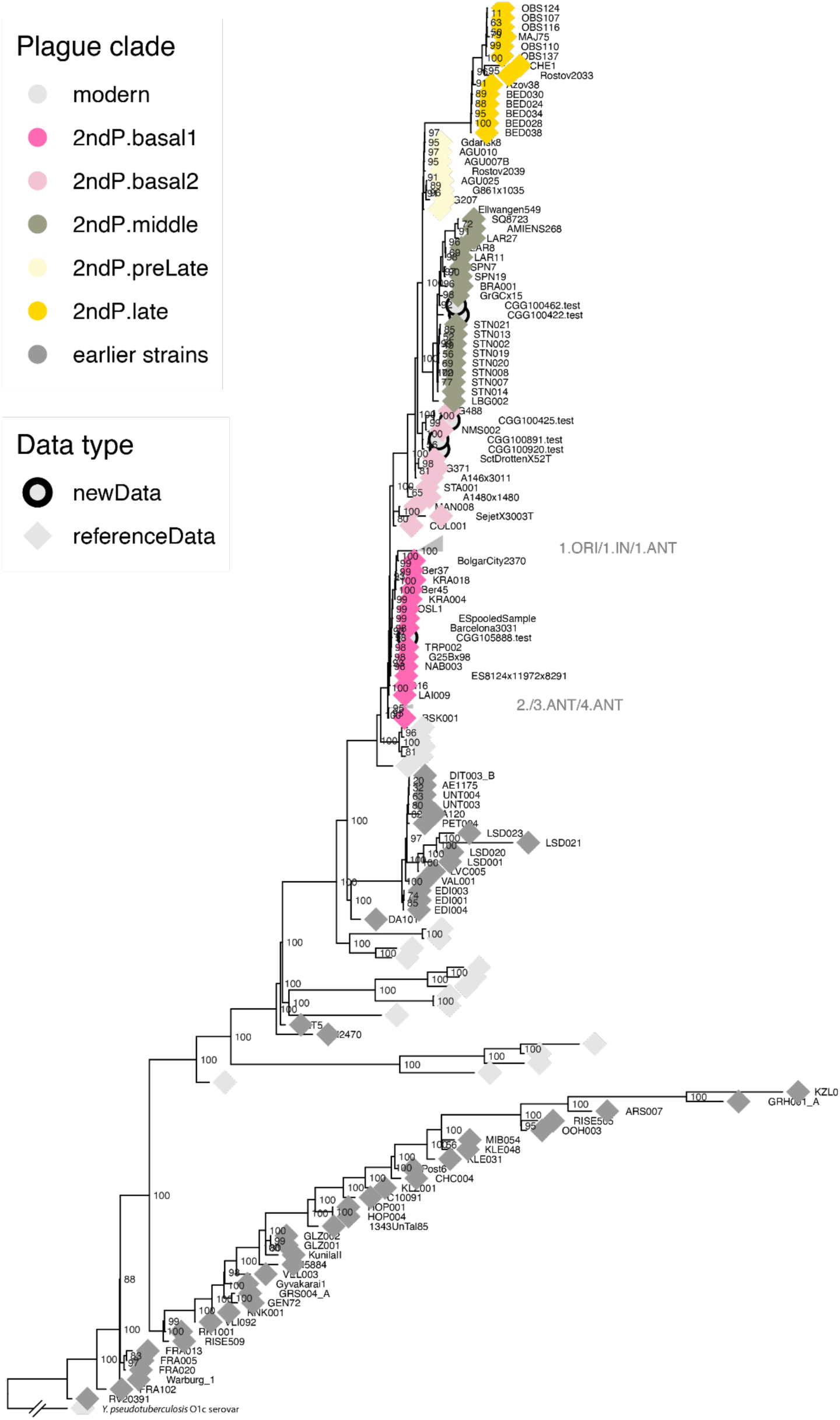
Full phylogenetic tree.

**Supplementary Figure 2.**
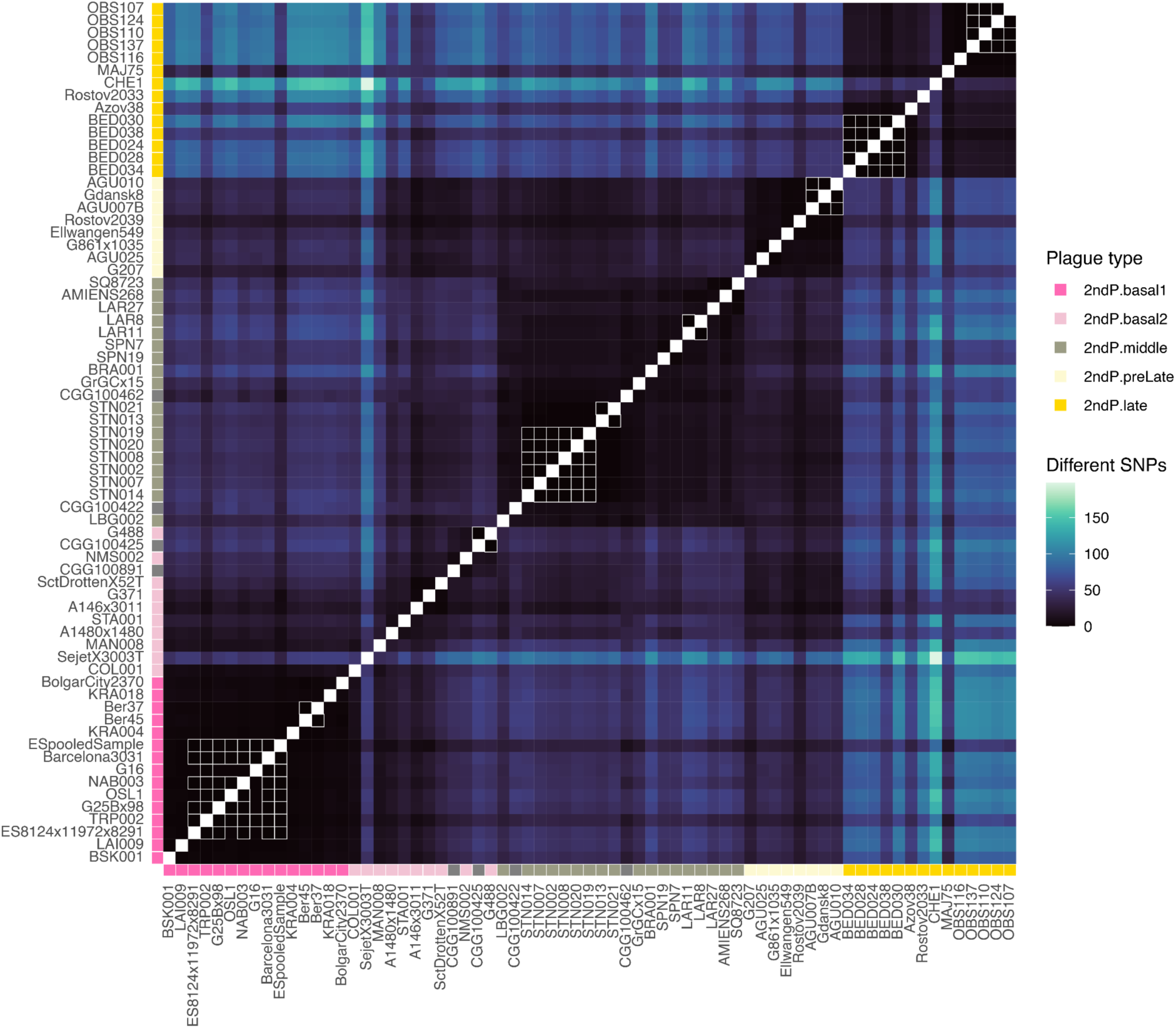
Heatmap of SNP differences across the plague genome for 2nd pandemic samples. Identical sample pairs that do not differ on any positions are highlighted with a white square.

**Supplementary Figure 3.**
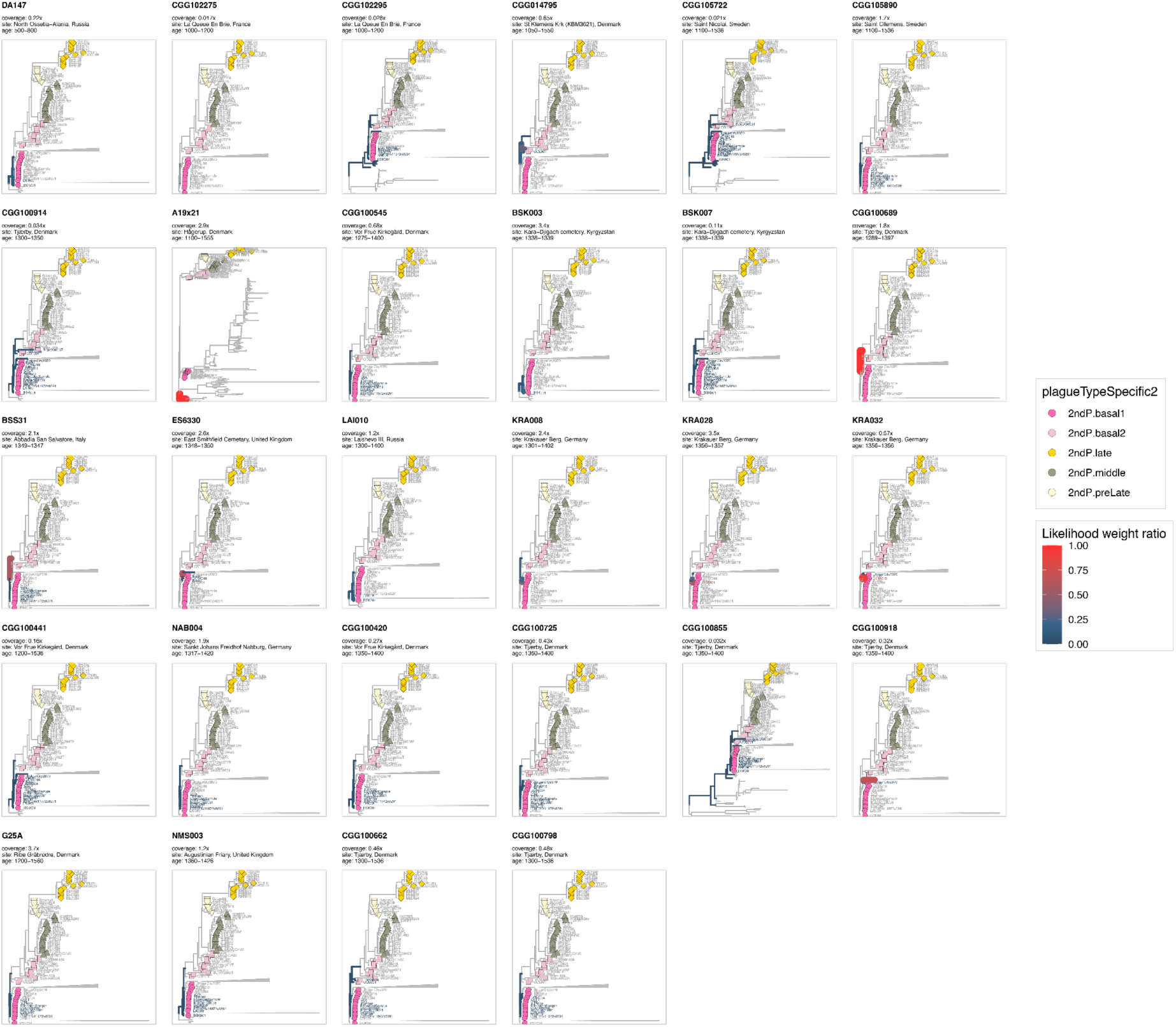
Placements of samples assigned to clade ‘basal1’.

**Supplementary Figure 4.**
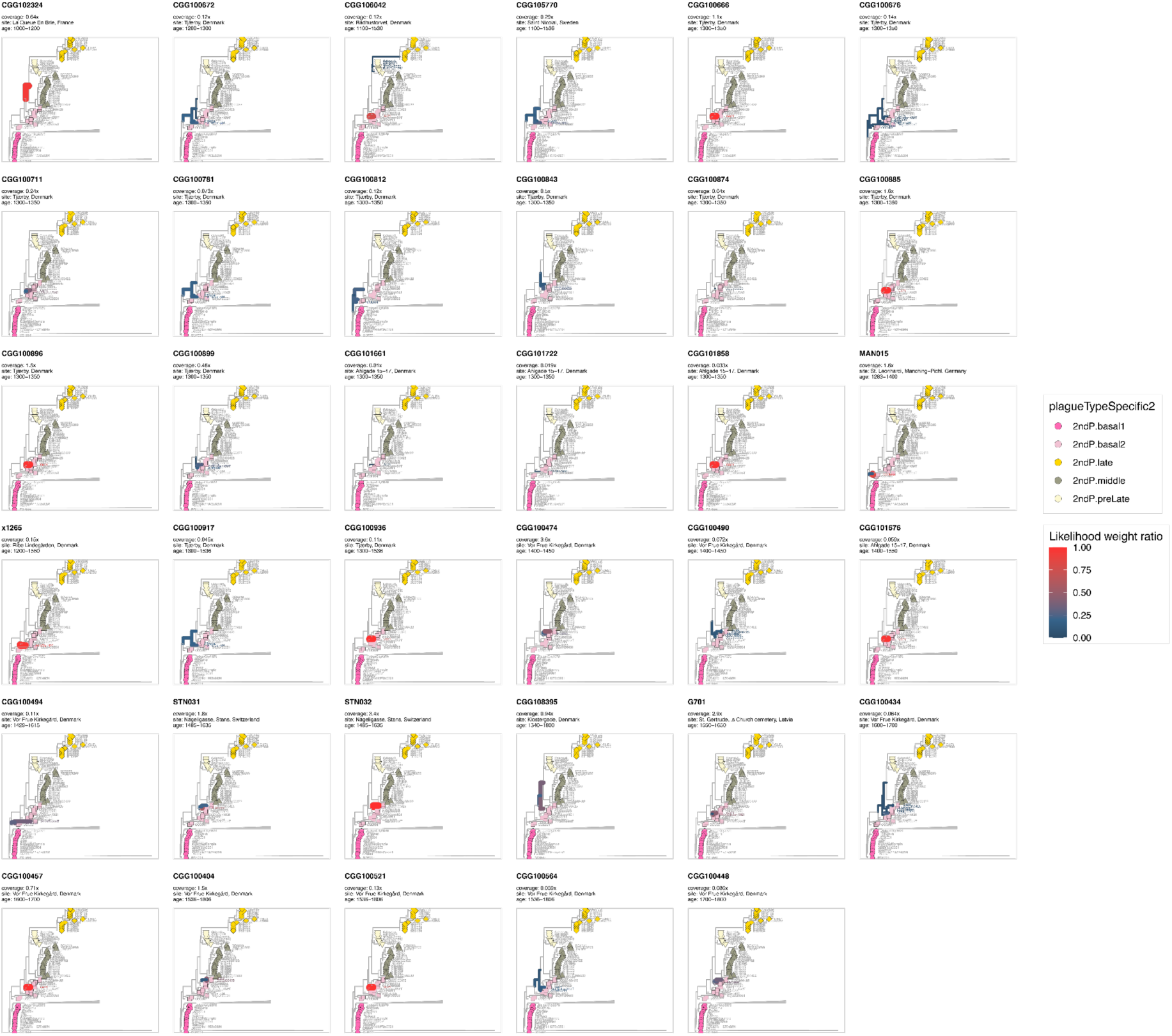
Placements of samples assigned to clade ‘basal2’.

**Supplementary Figure 5.**
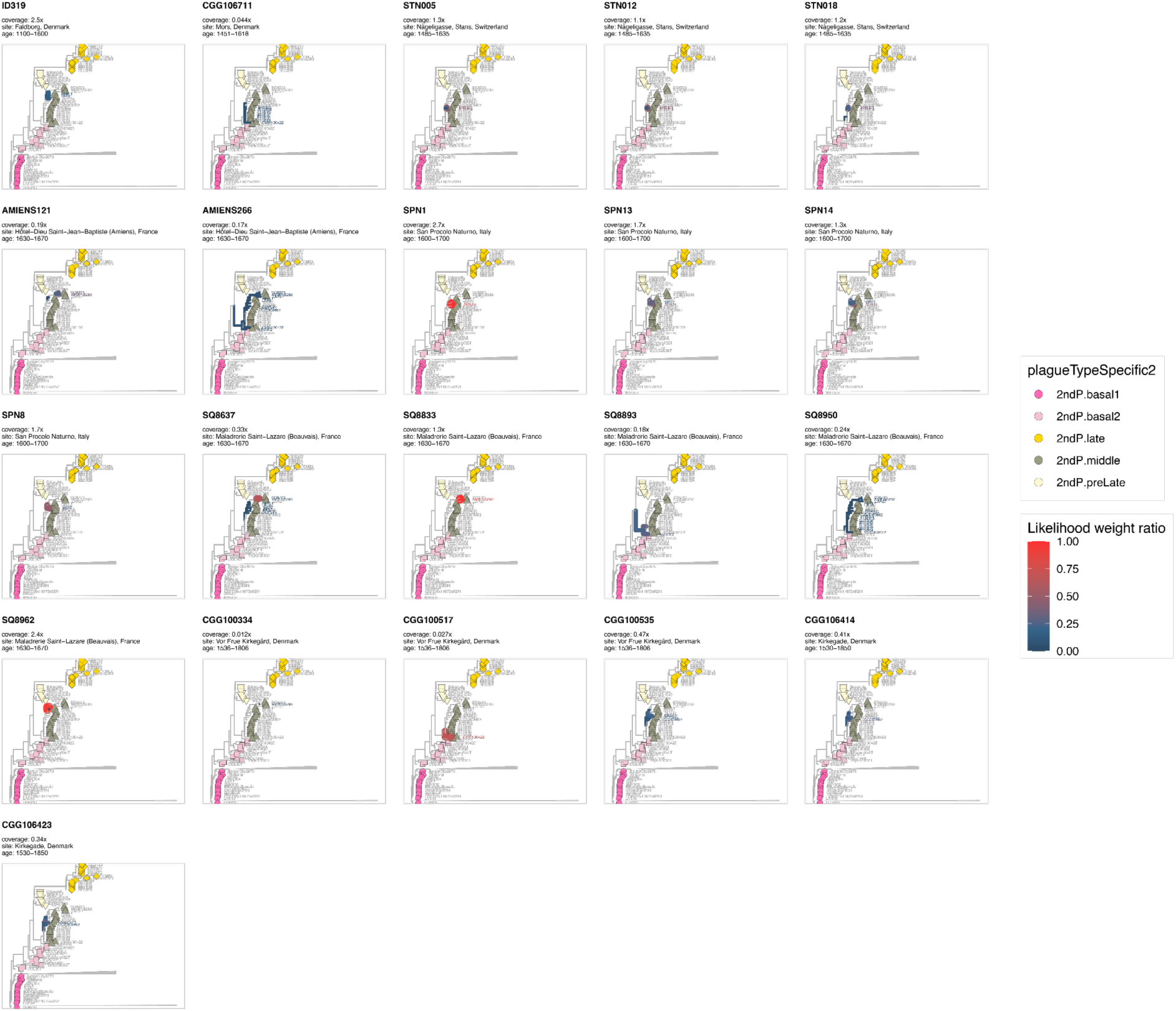
Placements of samples assigned to clade ‘middle’.

**Supplementary Figure 6.**
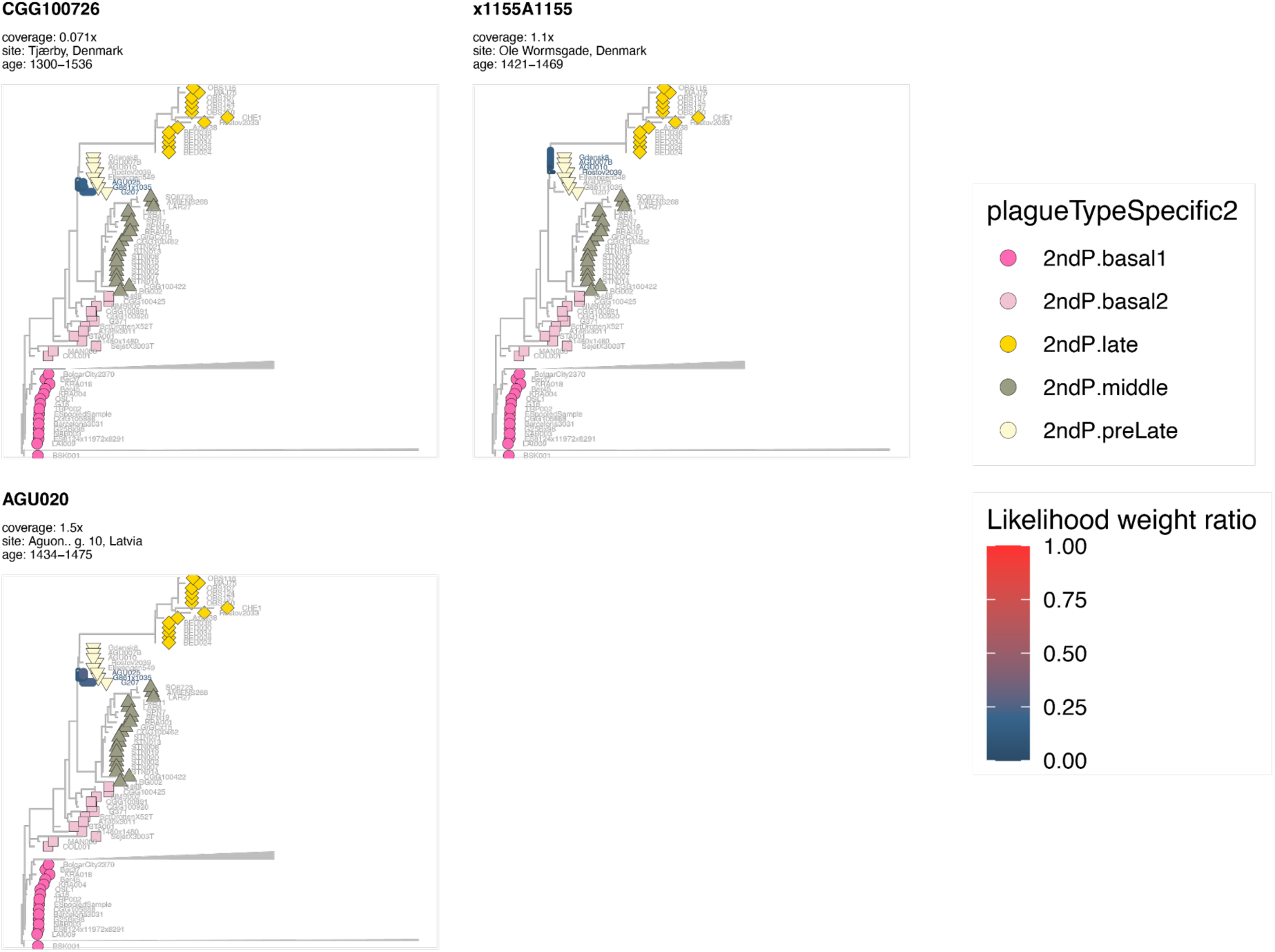
Placements of samples assigned to clade ‘preLate’.

**Supplementary Figure 7.**
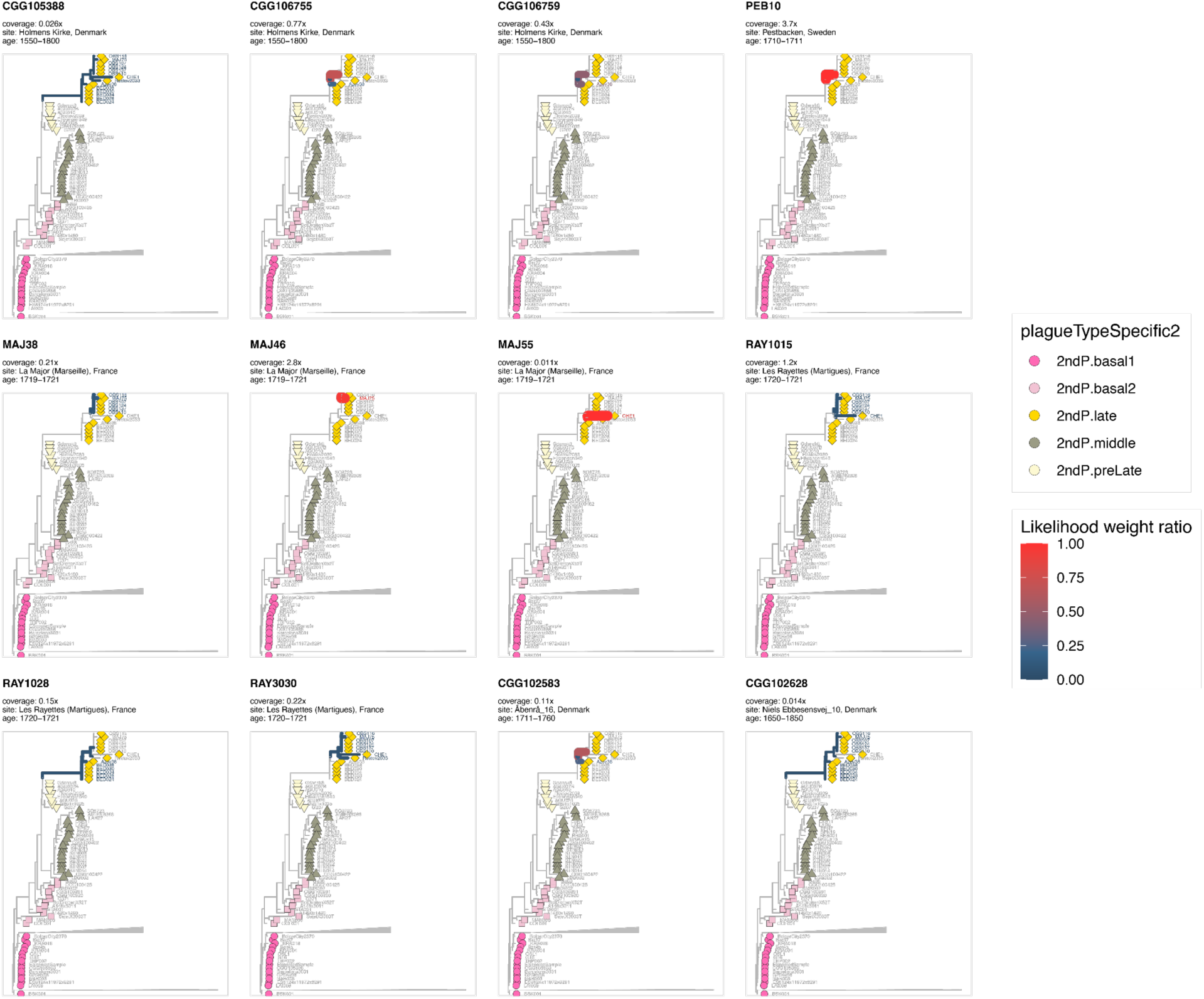
Placements of samples assigned to clade ‘late’.

**Supplementary Figure 8.**
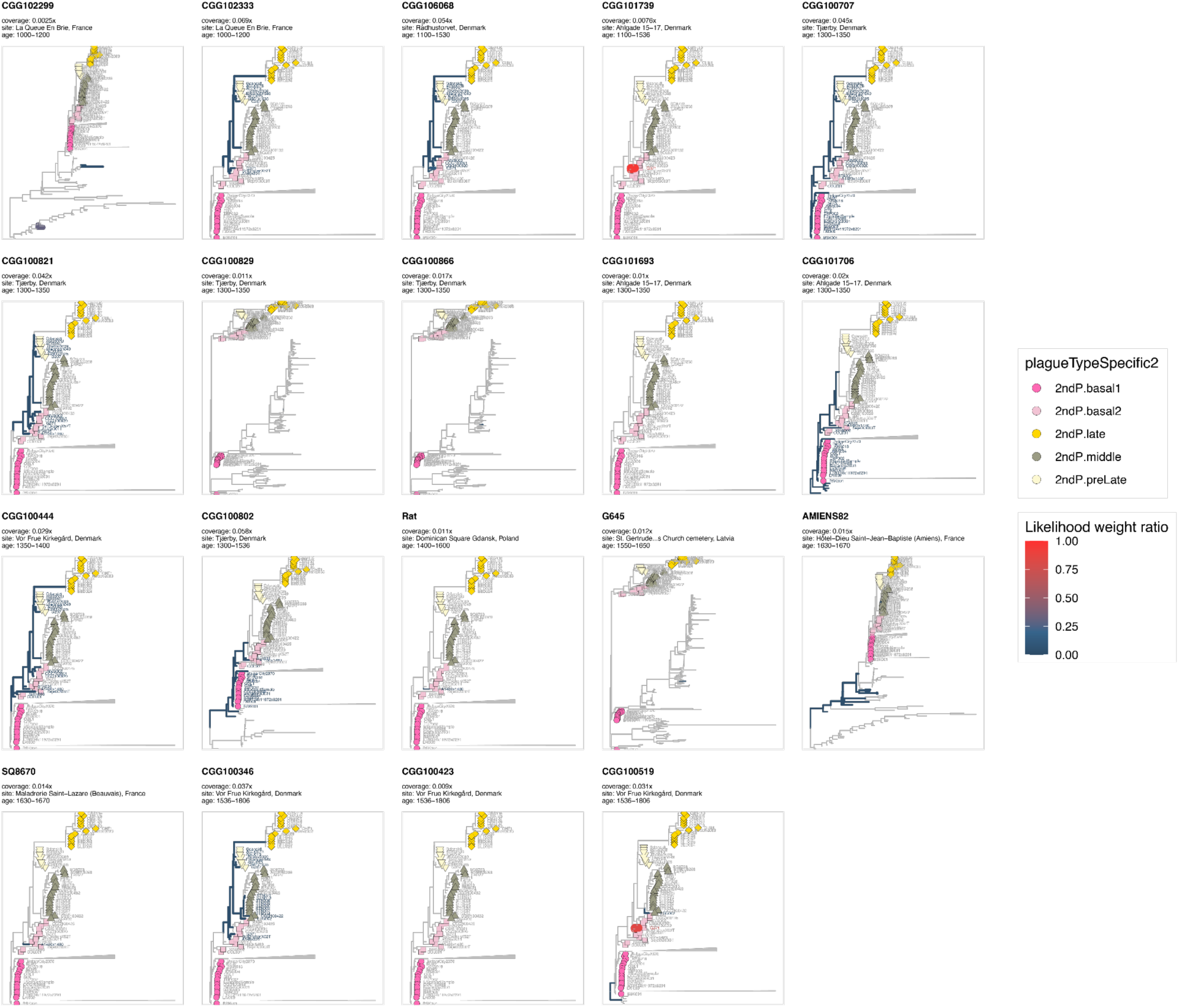
Placements of samples assigned to clade ‘lowcov’.

**Supplementary Figure 9.**
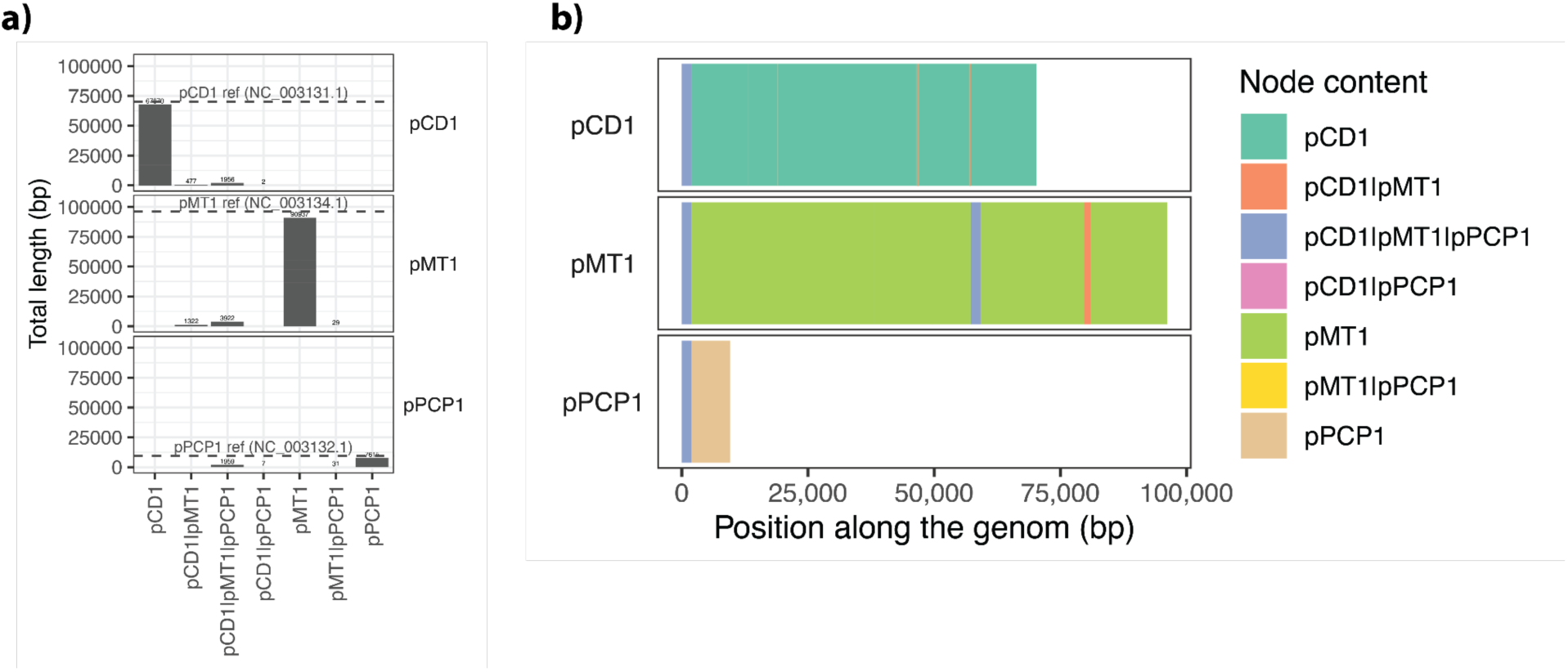
Overview of repeat regions in the canonical plague plasmids. **a)** This panel highlights that most of the length difference between the plasmid reference genomes (NC_003134.1, NC_003131.1 and NC_003132.1) and the length of the graph representing unique diversity from the three canonical Y. pestis plasmids are caused by varying numbers of repeat sequences, including insertion elements (IS100 (all plasmids), IS1541 (pMT1), and an IS285-like partial element (pMT1 and pCD1)). **b)** The three canonical Y. pestis plasmids have distinct evolutionary origins but they share mobile elements, most noticeable IS100 (blue), which is found once in pCD1 and pPCP1 and twice in pMT1. There are several overlaps between pMT1 and pCD1 (orange) including IS285-derived partial elements (PMID: 9748454). These homologous regions are often repetitive regions that could mediate plasmid rearrangements, which has been documented in one case (PMID: 21197443). pCD1 is the *Y. pestis* variant of the ancient pYV virulence plasmid shared between *Y. pseudotuberculosis* and *Y. enterocolitica*^82^. In contrast, pMT1 and pPCP1 were acquired during the emergence of *Y. pestis*. Following the LNBA-/LNBA+ split ∼5000 years BP, the ancestral LNBA+ lineage acquired the 19 kb *ymt* gene on pMT1 together with other flea-adaptive changes. The earliest sampled Y. pestis *ymt*+ variant is ∼3800 years old^22^. The acquisition of the *ymt* gene coincides with our observed increase in pMT1 copy number (Fig. 3). pPCP1 in the earliest *Y. pestis* lineages carried an ancestral *pla* gene variant, including *pla*-I259, and successive mutations accumulated during plague evolution. A V51L substitution was selected for on the ancestral LNBA+ lineage (coinciding with our observed sharp increase in pPCP1 copy number, Fig. 3)^83^, later an I259T substitution increased plasminogen activation and facilitated systemic dissemination, and the T259 variant was the most frequent variant in plague before the three historical pandemics^21^. Predominately following the Black Death, many plague samples show depletion of *pla* on multicopy pPCP1. This included recombination-mediated excision of *pla* from pPCP1 and integration into pCD1^13,20^; because pPCP1 is a high-copy plasmid and pCD1 a low copy plasmid, this resulted in a reduction of pla copy number and Pla proteolytic activity.

**Supplementary Figure 10.**
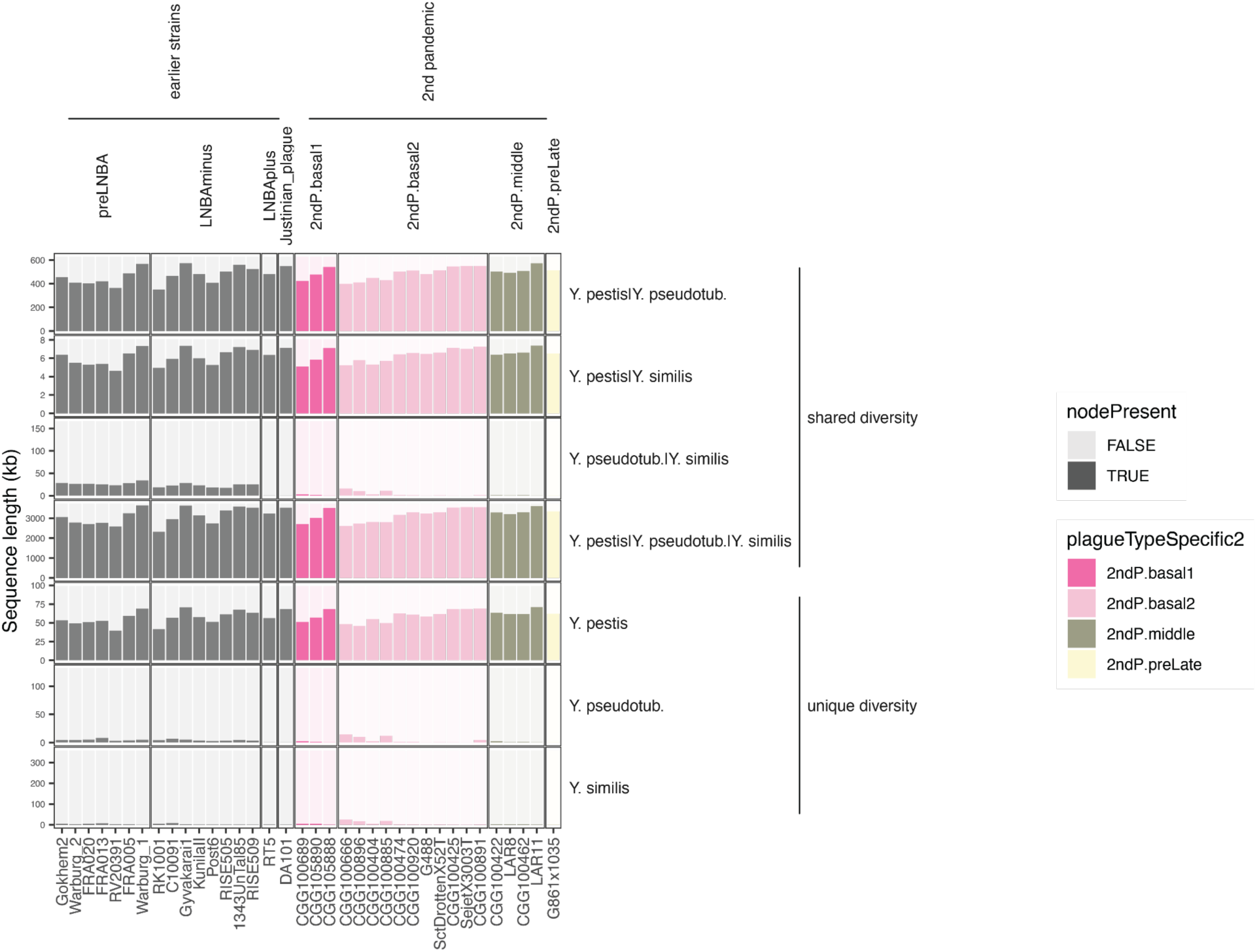
Variation graph-based chromosomal coverage. Vertically, the plot is stratified by diversity present in a single species (Y.pestis, Y.pseudotuberculosis or Y. similis) or shared among multiple species (shared diversity). Horizontally the plot is stratified by plague type. Note the variation in the scale of the y-axis.

**Supplementary Figure 11.**
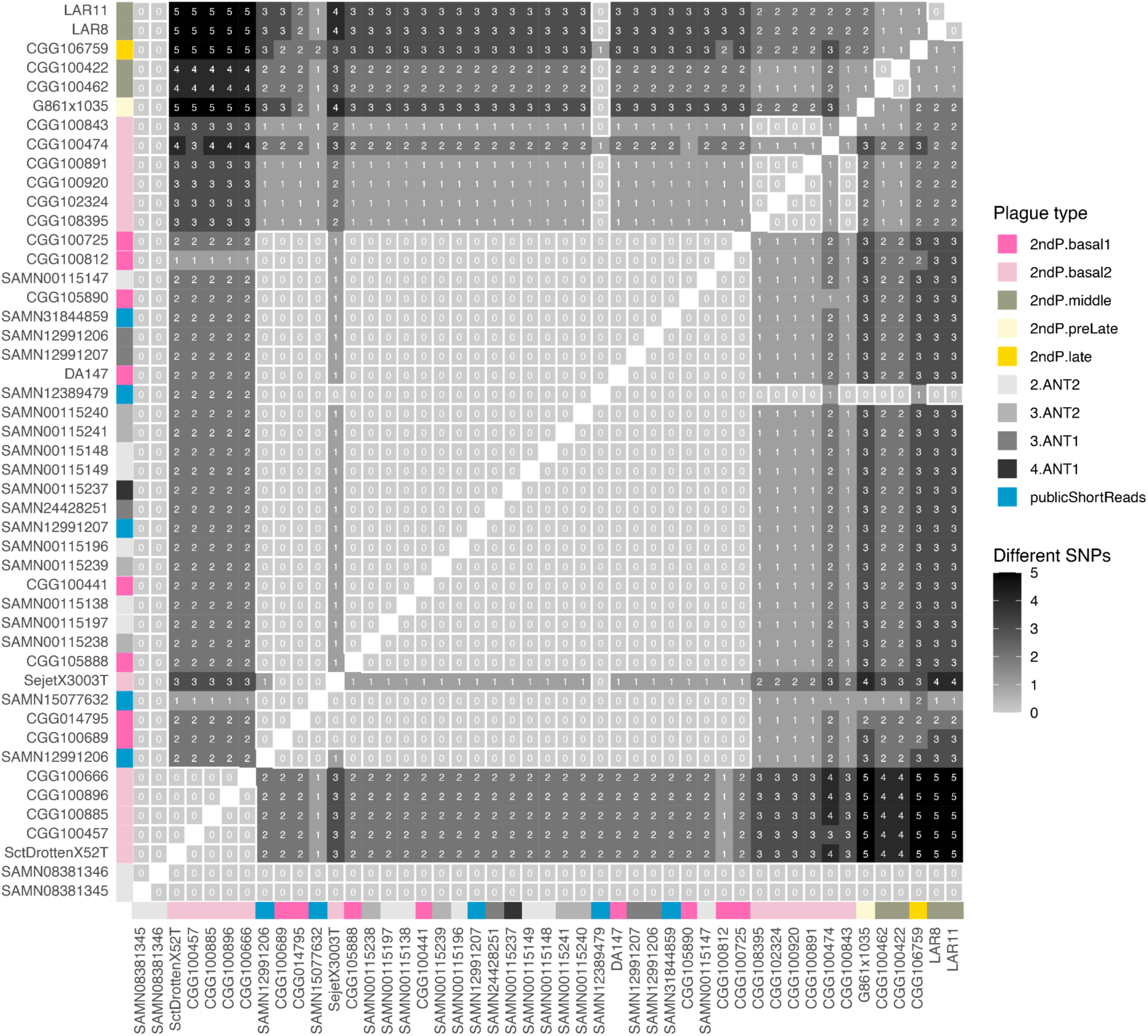
pTP33 heatmap of pairwise SNP differences. Identical sample pairs that do not differ on any positions are highlighted with a red square.

**Supplementary Figure 12.**
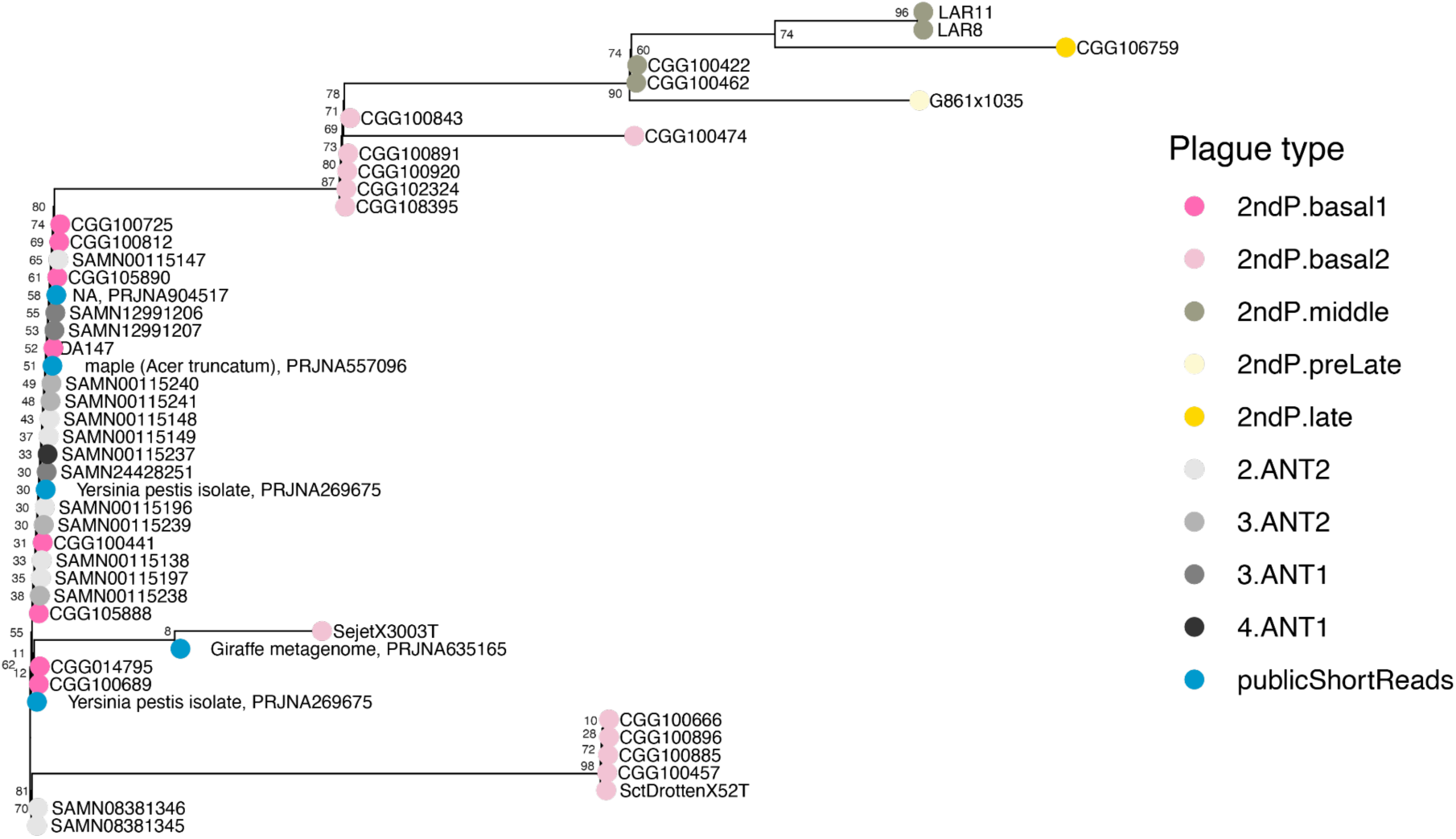
pTP33 phylogenetic tree.

**Supplementary Figure 13.**
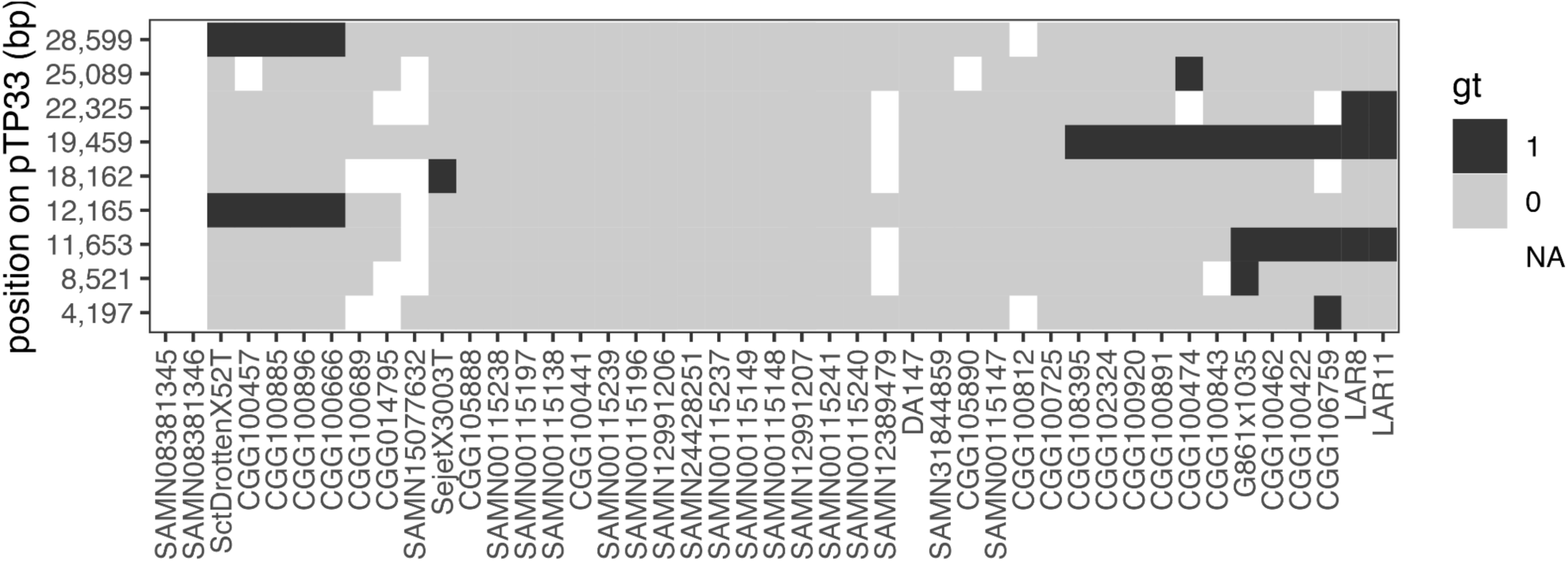
Variable positions from short read data mapped to pTP33.

**Supplementary Figure 14.**
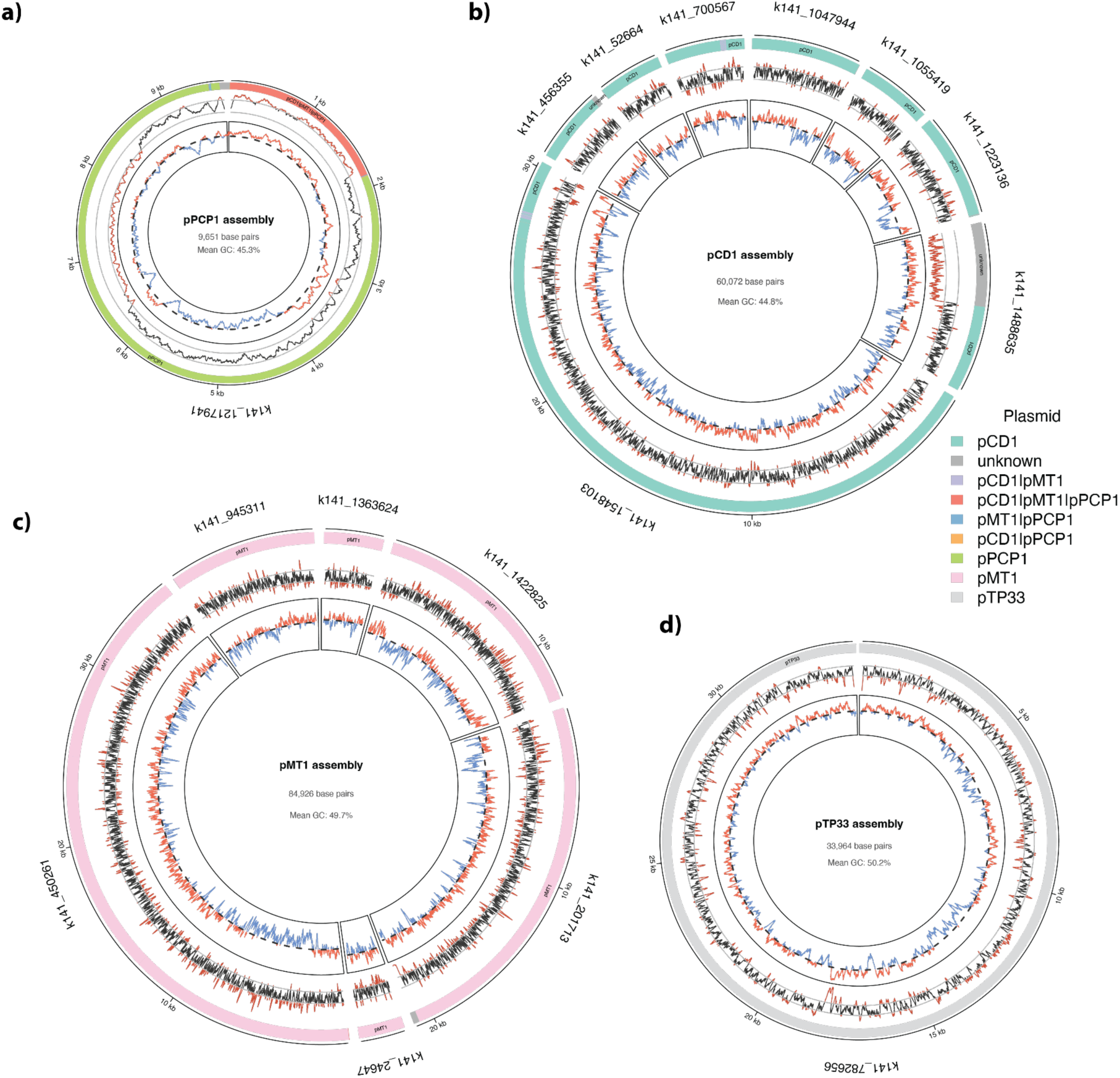
Assembly of plague plasmids from CGG100891. Outer circle with colored bars represents the category that each segment is assigned to based on the pangenome variation graph. The middle circle depicts the depth of coverage based on short reads mapped back to the assembly. Inner circler illustrates GC content across the plasmid at each position using a sliding window of 100bp. The stippled line represents the average GC content for the entire assembly (50.2%) while blue and red lines indicate GC content above and below the average respectively.

**Supplementary Figure 15.**
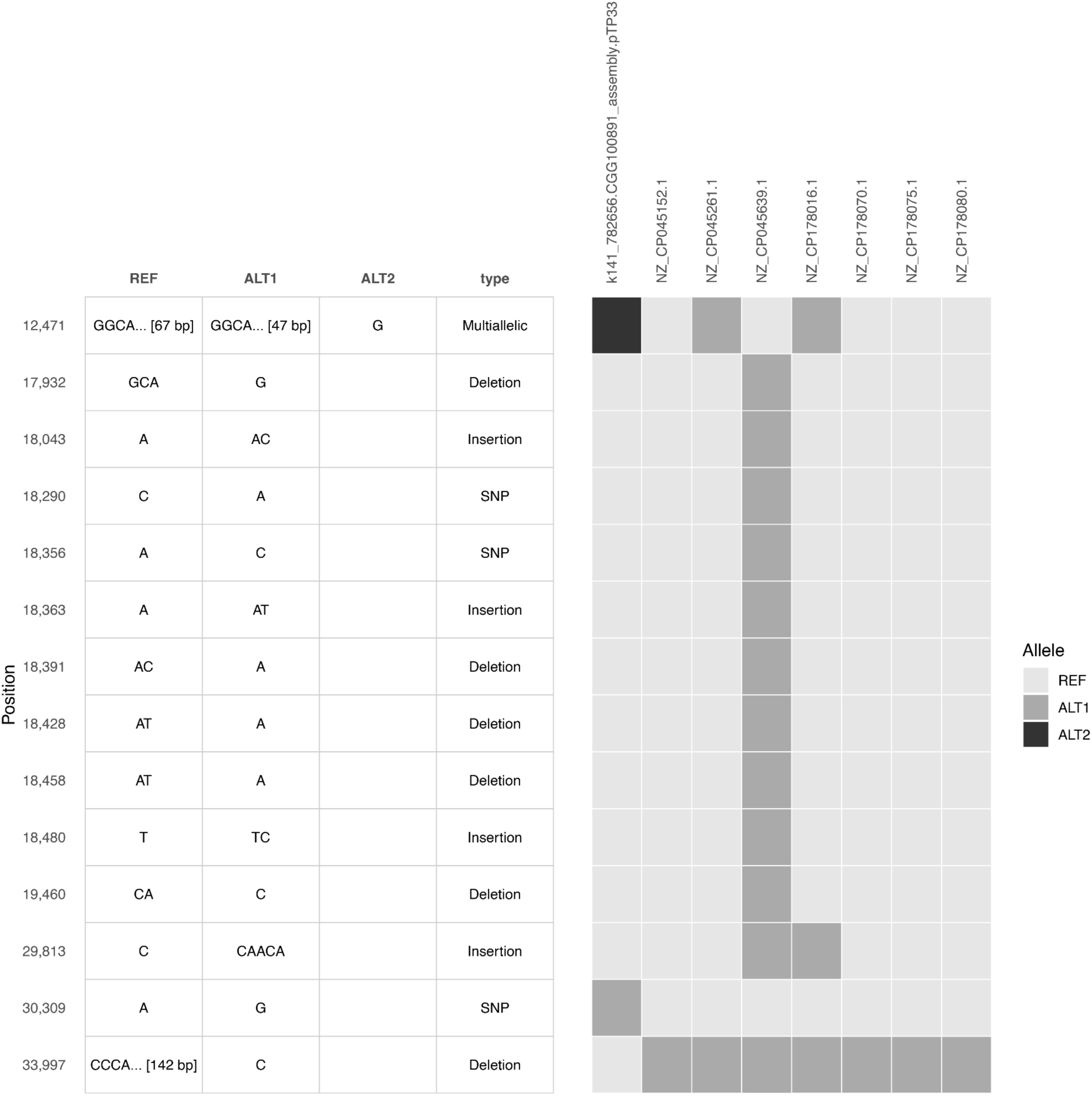
Variation among pTP33 assemblies. Left side panel contains information on each variant, while the right side panel depicts variants present in each of the 8 assemblies (7 publicly available assemblies and one generated for this study - K141_782656). As depicted, K141_782656 diverges from the consensus sequence at three positions (12 471, 30 309 and 33 997). The first of these (12,471) represents a deletion in a region with low complexity DNA, the second (30,309) represents an A->G single nucleotide polymorphism, and the last one (33,997) represents a duplication of a 142bp element at the very end of the assembly. Of these only the SNP at position 30,309 represents reliable evidence of genetic variation.

**Supplementary Figure 16.**
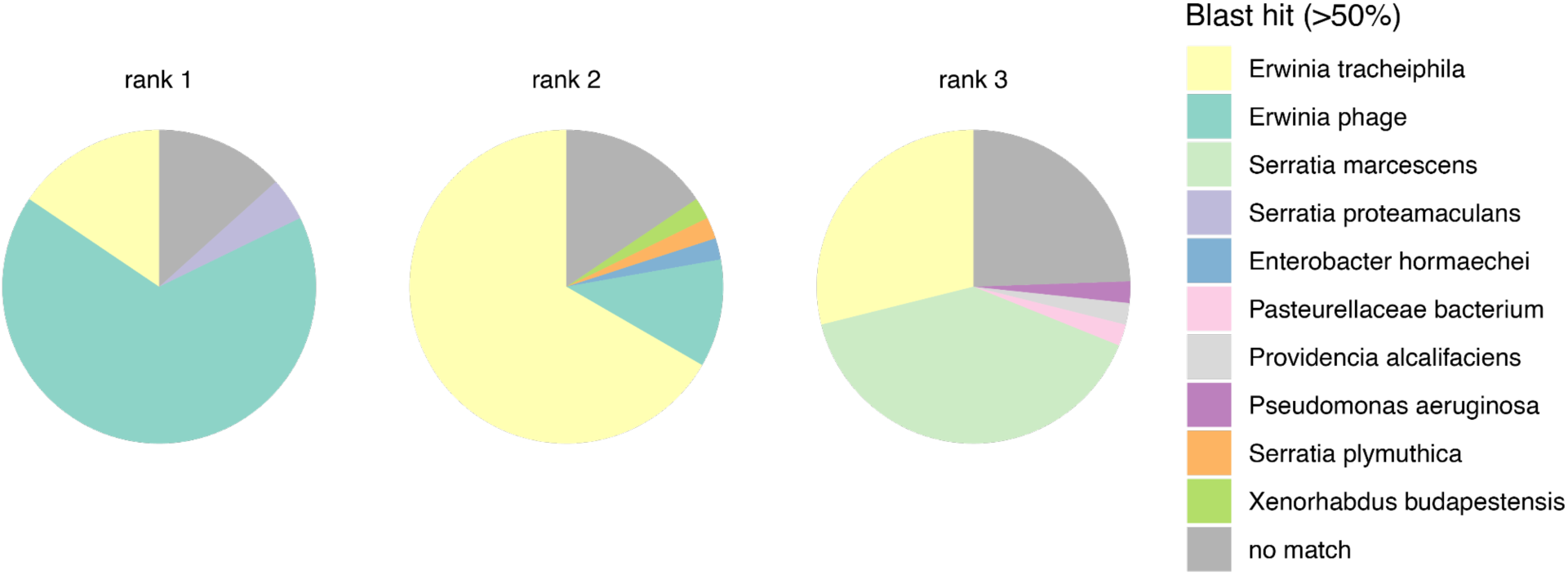
Best Blast hits for all CDS’es of pTP33.

**Supplementary Figure 17.**
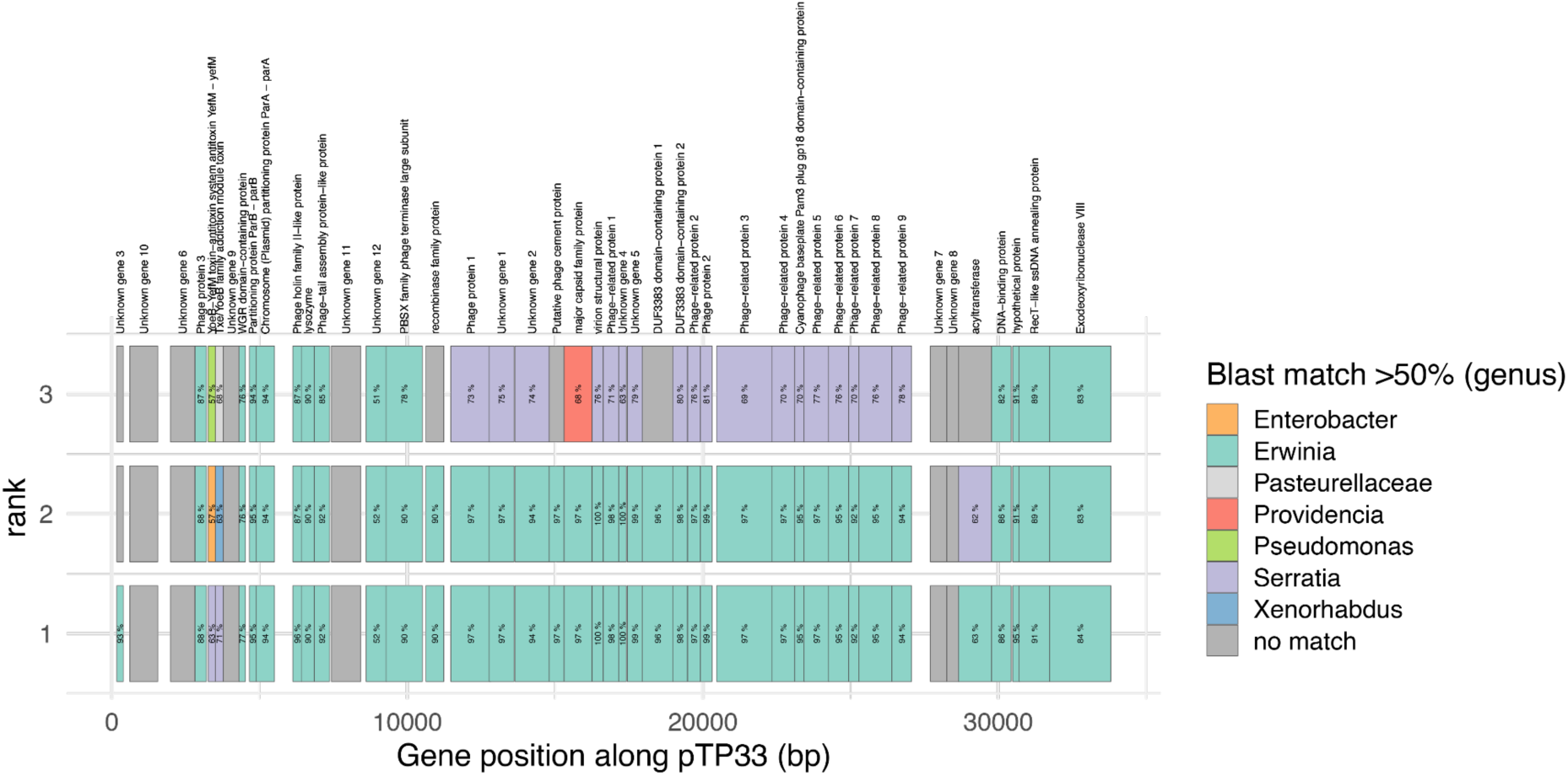
Best Blast hits for all CDS’es of pTP33 plotted along the plasmid.

**Supplementary Figure 18.**
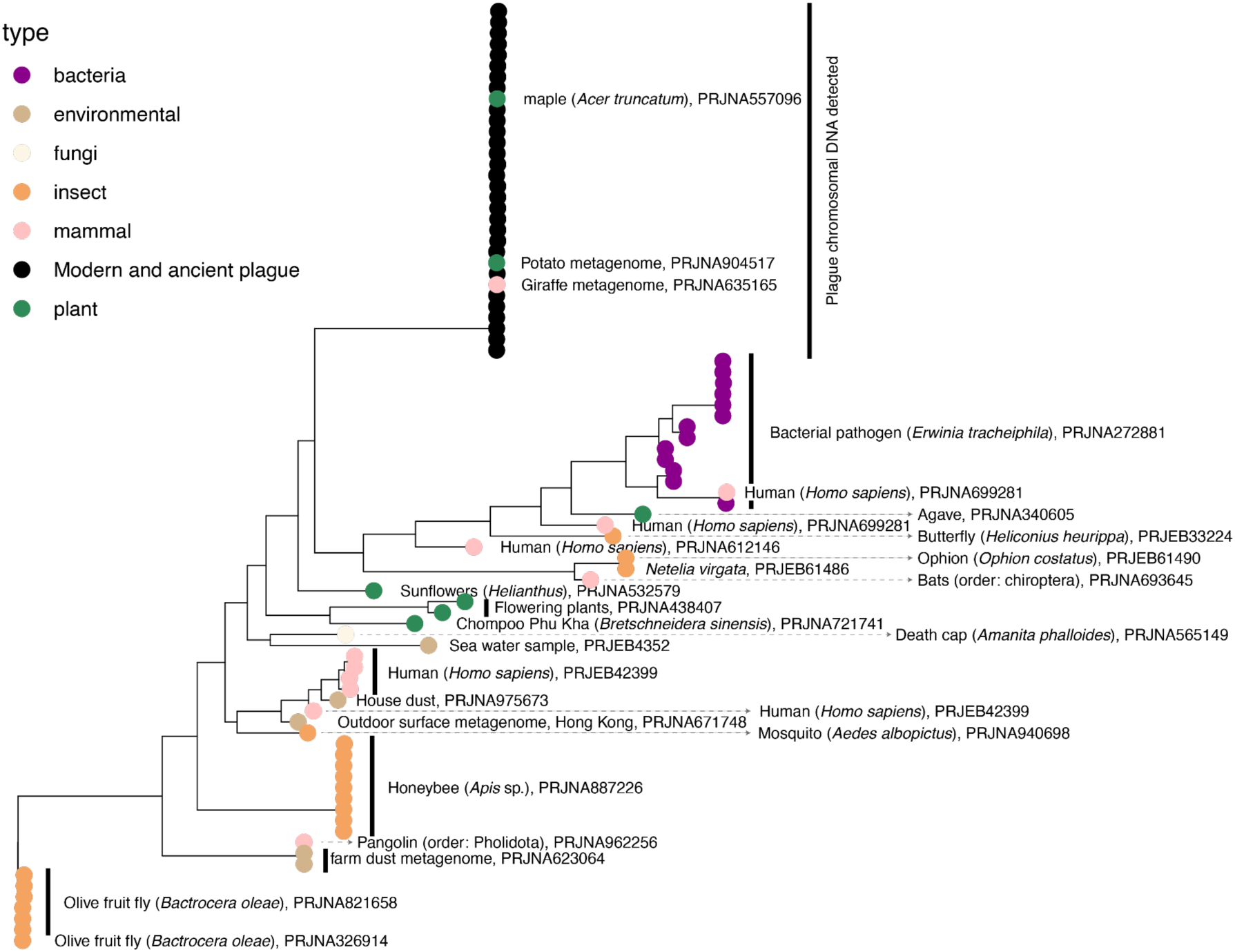
Logan results. Phylogenetic tree of pTP33 homologs from public short read metagenomes.

**Supplementary Figure 19.**
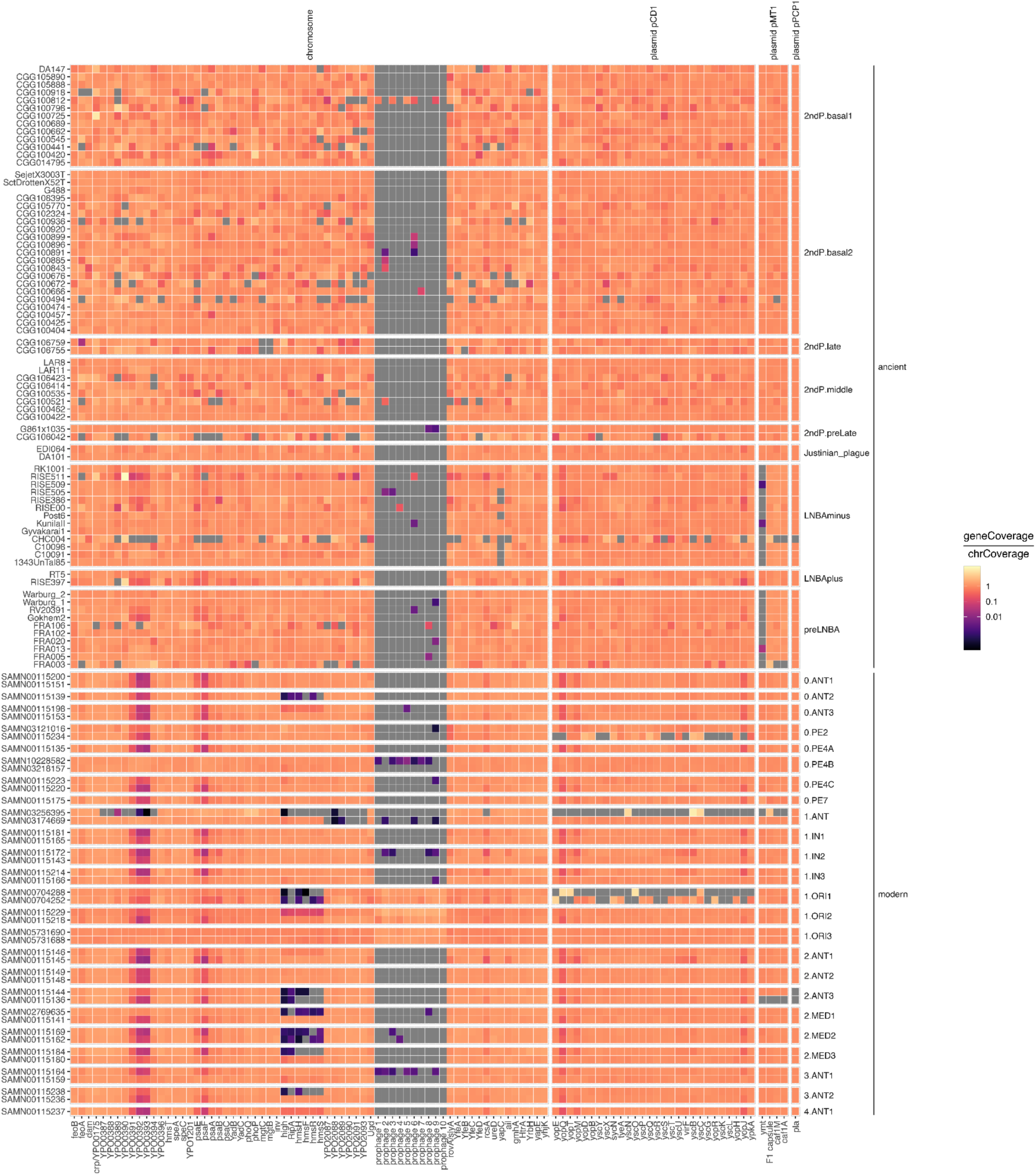
Virulence gene presence/absence.

**Supplementary Figure 20.**
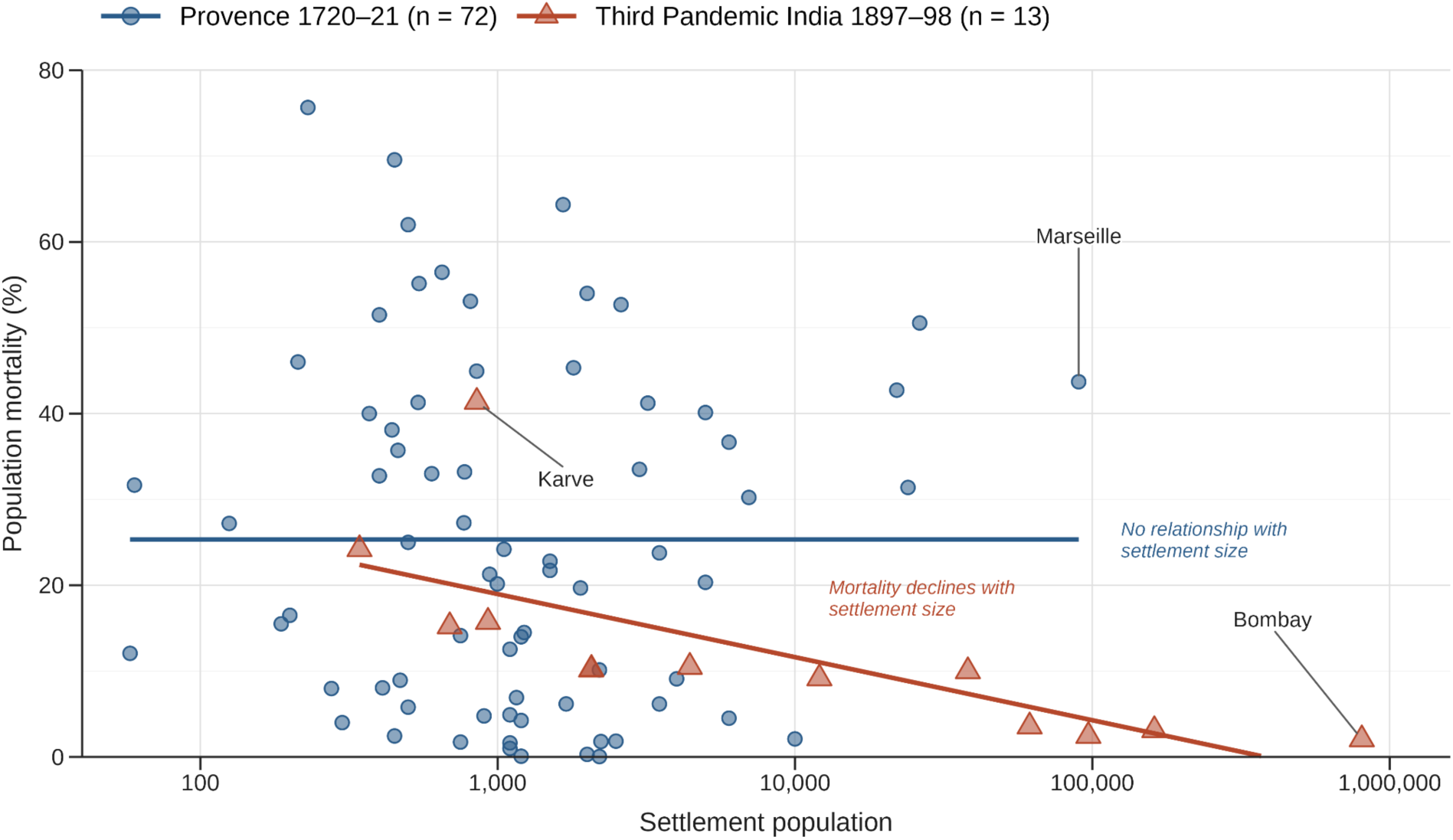
Settlement size and plague mortality during the Second and Third Plague Pandemics. Population mortality plotted against settlement population on a logarithmic x-axis. Second-Pandemic data comprise 72 localities affected during the Provence epidemic of 1720–21 (Biraben, 1968); Third-Pandemic data comprise 13 settlements in the Bombay Presidency in 1897–98 (Condon, 1900). Mortality was calculated from reported plague deaths and population. Lines show simple linear fits of mortality percentage against log10 population. Mortality was independent of settlement size in Provence but declined with increasing settlement size during the Third Pandemic.

## References

1. Spyrou, M. A. et al. The source of the Black Death in fourteenth-century central Eurasia. Nature 606, 718–724 (2022).

2. Benedictow, O. J. The Complete History of the Black Death. (Boydell Press, Woodbridge, England, 2024).

3. Sutherland, I. & Biraben, J.-N. Les Hommes et la Peste en France et dans les pays Europeens et Mediterraneens. Popul. Stud. (Camb.) 31, 603 (1977).

4. van Hoof, T. B., Bunnik, F. P. M., Waucomont, J. G. M., Kürschner, W. M. & Visscher, H. Forest re-growth on medieval farmland after the Black Death pandemic-Implications for atmospheric CO2 levels. Palaeogeogr. Palaeoclimatol. Palaeoecol. 237, 396–409 (2006).

5. DeWitte, S. N. Mortality risk and survival in the aftermath of the medieval Black Death. PLoS One 9, e96513 (2014).

6. Klunk, J. et al. Evolution of immune genes is associated with the Black Death. Nature 611, 312–319 (2022).

7. Schuenemann, V. J. et al. Targeted enrichment of ancient pathogens yielding the pPCP1 plasmid of Yersinia pestis from victims of the Black Death. Proc. Natl. Acad. Sci. U. S. A. 108, E746–52 (2011).

8. Lagerås, P. et al. Abandonment, agricultural change and ecology. in Environment, Society and the Black Death 30–68 (Oxbow Books, 2016).

9. Drancourt, M., Aboudharam, G., Signoli, M., Dutour, O. & Raoult, D. Detection of 400-year-old Yersinia pestis DNA in human dental pulp: an approach to the diagnosis of ancient septicemia. Proc. Natl. Acad. Sci. U. S. A. 95, 12637–12640 (1998).

10. Bos, K. I. et al. A draft genome of Yersinia pestis from victims of the Black Death. Nature 478, 506– 510 (2011).

11. Spyrou, M. A. et al. Phylogeography of the second plague pandemic revealed through analysis of historical Yersinia pestis genomes. Nat. Commun. 10, 4470 (2019).

12. Anisimov, A. P., Lindler, L. E. & Pier, G. B. Intraspecific diversity of Yersinia pestis. Clin. Microbiol. Rev. 17, 695–695 (2004).

13. Sidhu, R. K. et al. Attenuation of virulence in Yersinia pestis across three plague pandemics. Science 388, eadt3880 (2025).

14. Hinnebusch, B. J. et al. Role of Yersinia murine toxin in survival of Yersinia pestis in the midgut of the flea vector. Science 296, 733–735 (2002).

15. Yang, R. et al. Yersinia pestis and Plague: some knowns and unknowns. Zoonoses (Burlingt.) 3, 5 (2023).

16. Ermoli, F., Spahn, C., Sadhir, I., Bode, H. B. & Diepold, A. Activation of the Yersinia type III secretion system induces large-scale spatial reorganization of chromosomal and virulence plasmid DNA. Cell Rep. 44, 116592 (2025).

17. Rasmussen, S. et al. Early divergent strains of Yersinia pestis in Eurasia 5,000 years ago. Cell 163, 571–582 (2015).

18. Rodríguez-Beltrán, J., DelaFuente, J., León-Sampedro, R., MacLean, R. C. & San Millán, Á. Beyond horizontal gene transfer: the role of plasmids in bacterial evolution. Nat. Rev. Microbiol. 19, 347–359 (2021).

19. Manning, S. W., Birch, J., Conger, M. A. & Sanft, S. Resolving time among non-stratified short-duration contexts on a radiocarbon plateau: Possibilities and challenges from the ad 1480–1630 example and northeastern North America. Radiocarbon 62, 1785–1807 (2020).

20. Susat, J. et al. Yersinia pestis strains from Latvia show depletion of the pla virulence gene at the end of the second plague pandemic. Sci. Rep. 10, 14628 (2020).

21. Zimbler, D. L., Schroeder, J. A., Eddy, J. L. & Lathem, W. W. Early emergence of Yersinia pestis as a severe respiratory pathogen. Nat. Commun. 6, 7487 (2015).

22. Spyrou, M. A. et al. Analysis of 3800-year-old Yersinia pestis genomes suggests Bronze Age origin for bubonic plague. Nat. Commun. 9, 2234 (2018).

23. Dai, R., et al. A novel mechanism of streptomycin resistance in Yersinia pestis: Mutation in the rpsL gene. PLoS Negl. Trop. Dis. 15, e0009324 (2021).

24. Afanas’ev, M. V. et al. Analysis of complete sequence of cryptic plasmid pTP33 from Yersinia pestis isolated in Tuva natural focus of plague. Genetika 52, 1012–1020 (2016).

25. Bazanova, L. P., Tokmakova, E. G., Voronova, G. A. & Balakhonov, S. V. Influence of Yersinia pestis Plasmid content on biofilm formation in fleas with different vector activity. Journal of microbiology, epidemiology and immunobiology 95, 76–83 (2018).

26. Chikhi, R. et al. Logan: Planetary-scale genome assembly surveys life’s diversity. bioRxivorg (2025) doi:10.1101/2024.07.30.605881.

27. Shapiro, L. R. et al. Draft genome sequence of Erwinia tracheiphila, an economically important bacterial pathogen of cucurbits. Genome Announc. 3, e00482–15 (2015).

28. Bramanti, B., Dean, K. R., Walløe, L. & Chr Stenseth, N. The third plague pandemic in Europe. Proc. Biol. Sci. 286, 20182429 (2019).

29. Andrianaivoarimanana, V., et al. Understanding the persistence of plague foci in Madagascar. PLoS Negl. Trop. Dis. 7, e2382 (2013).

30. Harimalala, M. Current knowledge on fleas (Siphonaptera) associated with human plague transmission in Madagascar. J. Med. Entomol. 62, 749–759 (2025).

31. Bazanova, L. P., Tokmakova, E. G., Voronova, G. A., Balakhonov, S. V. & Innokent’eva, T. I. Peculiarities of interaction between Citellophilus tesquorum altaicus (ioff, 1936) flea and Yersinia pestis with various Plasmid composition. Probl. Osobo Opas. Infekc. 41–44 (2017).

32. Bazanova, L. P., Nikitin, A. Y. & Tokmakova, E. G. The Yersinia pestis aggregation as a functional adaptation to the flea organism (review). Journal of microbiology, epidemiology and immunobiology 100, 240–250 (2023).

33. He, Z. et al. Distribution and characteristics of human plague cases and Yersinia pestis isolates from 4 Marmota plague foci, China, 1950-2019. Emerg. Infect. Dis. 27, 2544–2553 (2021).

34. Medvedev, S. G., Kotti, B. K. & Verzhutsky, D. B. Diversity of fleas (Siphonaptera), vectors of plague pathogens: The flea Citellophilus tesquorum (Wagner, 1898), a parasite of ground squirrels of the genus Spermophilus. Entomol. Rev. 99, 565–579 (2019).

35. Ingrassia, G. F. Informatione del pestifero, et contagioso morbo. (Giovan Matteo Mayda, Palermo, 1576).

36. Rodenwaldt, E. Pest in Venedig 1575--1577: Ein Beitrag zur Frage der Infektkette bei den Pestepidemien West-Europas. (Springer, Berlin and Heidelberg, 1953).

37. Ó Glacáin, N. *T*ractatus de peste, seu, Brevis, facilis, & experta methodus curandi pestem. (Typis Raymundi Colomerii, Tolosae [Toulouse], 1629).

38. Dean, K. R. et al. Human ectoparasites and the spread of plague in Europe during the Second Pandemic. Proc. Natl. Acad. Sci. U. S. A. 115, 1304–1309 (2018).

39. Barbieri, R. et al. Differential word expression analyses highlight plague dynamics during the second pandemic. R. Soc. Open Sci. 9, 210039 (2022).

40. Egilssonar, J. Biskupa-Annálar Jóns Egilssonar. in Safn Til Sögu Íslands Og Íslenzkra Bókmenta Að Fornu Og Nýju (ed. Sigurðsson, J.) vol. 1 15–136 (Hið íslenzka bókmenntafélag, Kaupmannahöfn, 1856).

41. Ferguson, C. R. et al. The Genomic Landscape of Post-Black Death Epidemics in Northern Europe and the Caucasus. bioRxiv : the preprint server for biology https://doi.org/10.64898/2026.08.25.746822 (2026) doi:10.64898/2026.08.25.746822.

42. Kjær, K. et al. Environmental DNA reveals Reykjavík’s human and ecological history. bioRxiv 2025.10.08.681091 (2025) doi:10.1101/2025.10.08.681091.

43. Karlsson, G. Plague without Rats: The Case of Fifteenth-Century Iceland. Journal of Medieval History 22, 263–284 (1996).

44. Drancourt, M., Houhamdi, L. & Raoult, D. Yersinia pestis as a telluric, human ectoparasite-borne organism. Lancet Infect. Dis. 6, 234–241 (2006).

45. Burroughs, A. L. Sylvatic Plague Studies: The Vector Efficiency of Nine Species of Fleas Compared with Xenopsylla Cheopis. Journal of Hygiene 45, 371–396 (1947).

46. Miarinjara, A., Bland, D. M., Belthoff, J. R. & Hinnebusch, B. J. Poor vector competence of the human flea, Pulex irritans, to transmit Yersinia pestis. Parasit. Vectors 14, 317 (2021).

47. Bland, D. M., Long, D., Rosenke, R. & Hinnebusch, B. J. Yersinia pestis can infect the Pawlowsky glands of human body lice and be transmitted by louse bite. PLoS Biol. 22, e3002625 (2024).

48. Sikora, M. et al. The spatiotemporal distribution of human pathogens in ancient Eurasia. Nature 643, 1011–1019 (2025).

49. Willerslev, E. & Cooper, A. Ancient DNA. Proc. Biol. Sci. 272, 3–16 (2005).

50. Damgaard, P. B. et al. Improving access to endogenous DNA in ancient bones and teeth. Sci. Rep. 5, 11184 (2015).

51. Rohland, N., Glocke, I., Aximu-Petri, A. & Meyer, M. Extraction of highly degraded DNA from ancient bones, teeth and sediments for high-throughput sequencing. Nat. Protoc. 13, 2447–2461 (2018).

52. McColl, H. et al. Steppe Ancestry in western Eurasia and the spread of the Germanic Languages. bioRxiv 2024.03.13.584607 (2024) doi:10.1101/2024.03.13.584607.

53. Meyer, M. & Kircher, M. Illumina sequencing library preparation for highly multiplexed target capture and sequencing. Cold Spring Harb. Protoc. 2010, db.prot5448 (2010).

54. A Fast and Efficient Single-Stranded Genomic Library Preparation Method Optimized for Ancient DNA Open Access.

55. Wagner, D. M. et al. Yersinia pestis and the plague of Justinian 541-543 AD: a genomic analysis. Lancet Infect. Dis. 14, 319–326 (2014).

56. Schubert, M., Lindgreen, S. & Orlando, L. AdapterRemoval v2: rapid adapter trimming, identification, and read merging. BMC Res. Notes 9, 88 (2016).

57. Langmead, B. & Salzberg, S. L. Fast gapped-read alignment with Bowtie 2. Nat. Methods 9, 357–359 (2012).

58. Li, H. et al. The Sequence Alignment/Map format and SAMtools. Bioinformatics 25, 2078–2079 (2009).

59. McKenna, A. et al. The Genome Analysis Toolkit: a MapReduce framework for analyzing next-generation DNA sequencing data. Genome Res. 20, 1297–1303 (2010).

60. Danecek, P. et al. Twelve years of SAMtools and BCFtools. Gigascience 10, giab008 (2021).

61. Yu, G., Lam, T. T.-Y., Zhu, H. & Guan, Y. Two methods for mapping and visualizing associated data on phylogeny using ggtree. Mol. Biol. Evol. 35, 3041–3043 (2018).

62. Yu, G., Smith, D. K., Zhu, H., Guan, Y. & Lam, T. T.-Y. Ggtree: An r package for visualization and annotation of phylogenetic trees with their covariates and other associated data. Methods Ecol. Evol. 8, 28–36 (2017).

63. Barbera, P. et al. EPA-ng: Massively parallel evolutionary placement of genetic sequences. Syst. Biol. 68, 365–369 (2019).

64. Garrison, E. et al. Building pangenome graphs. Nat. Methods https://doi.org/10.1038/s41592-024-02430-3 (2024) doi:10.1038/s41592-024-02430-3.

65. Sirén, J. et al. Pangenomics enables genotyping of known structural variants in 5202 diverse genomes. Science 374, abg8871 (2021).

66. Quinlan, A. R. & Hall, I. M. BEDTools: a flexible suite of utilities for comparing genomic features. Bioinformatics 26, 841–842 (2010).

67. Chen, S., Zhou, Y., Chen, Y. & Gu, J. fastp: an ultra-fast all-in-one FASTQ preprocessor. Bioinformatics 34, i884–i890 (2018).

68. Li, D., Liu, C.-M., Luo, R., Sadakane, K. & Lam, T.-W. MEGAHIT: an ultra-fast single-node solution for large and complex metagenomics assembly via succinct de Bruijn graph. Bioinformatics 31, 1674– 1676 (2015).

69. Hyldgard, I. M. Tjaerby Odekirke Og Kirkegard. (Aarhus University Press, 2016).

70. Society and the Black Death: An interdisciplinary approach to the late-medieval crisis in Sweden. Environment.

71. Grove, J. M. The Little Ice Age. (Routledge, London, England, 2004).

72. Biraben, J.-N. Certain Demographic Characteristics of the Plague Epidemic in France, 1720--22. Daedalus 97, 536–545 (1968).

73. Condon, J. K. The Bombay Plague: Being a History of the Progress of Plague in the Bombay Presidency from September 1896 to June 1899. (Education Society’s Steam Press, Bombay, 1900).

74. Jedwab, R., Johnson, N. D. & Koyama, M. The Economic Impact of the Black Death. Journal of Economic Literature 60, 132–178 (2022).

75. Greenwood, M. Statistical Investigation of Plague in the Punjab. Third Report: On Some of the Factors Which Influence the Prevalence of Plague. The Journal of Hygiene 11, 62–156 (1911).

76. Advisory Committee Appointed by the Secretary of State for India, the Royal Society, and the Lister Institute. XXIV. General Considerations Regarding the Spread of Infection, Infectivity of Houses, Etc. in Bombay City and Island. The Journal of Hygiene 7, 874–894 (1907).

77. Morello, C. L’arrivo Della Peste Dall’esterno. Preprint at (1584).

78. Palmer, R. J. The Control of Plague in Venice and Northern Italy, 1348--1600. (University of Kent at Canterbury, 1978). doi:10.22024/UniKent/01.02.86158.

79. O’Glacan, N. Tractatus de Peste, Seu, Brevis, Facilis, et Experta Methodus Curandi Pestem. (Raymundus Colomerius, Tolosa, 1629).

80. Tadino, A. Raguaglio Dell’origine et Giornali Successi Della Gran Peste Contagiosa, Venefica, & Malefica Seguita Nella Città Di Milano, & Suo Ducato Dall’anno 1629 Fino All’anno 1632. (Filippo Ghisolfi, Milano, 1648).

81. de Diemerbroeck, I. Tractatus de Peste. (Joannes Blaeu, Amsterdam, 1665).

82. Hu, P. et al. Structural organization of virulence-associated plasmids of Yersinia pestis. J. Bacteriol. 180, 5192–5202 (1998).

83. Sun, B. et al. A prehistoric East-Asian Yersinia pestis genome and a ∼5.3 ka trans-Eurasian expansion of plague. Nat. Commun. 17, (2026).

